# Optimization of molecular methods for detection and quantification of specific duckweed-bacteria interactions

**DOI:** 10.1101/2023.01.04.522651

**Authors:** Kenneth Acosta, Shawn Sorrels, William Chrisler, Weijuan Huang, Sarah Gilbert, Thomas Brinkman, Todd P. Michael, Sarah Lebeis, Eric Lam

## Abstract

Bacterial colonization dynamics of plants can differ between phylogenetically similar bacterial strains as well as in the context of complex bacterial communities. Quantitative studies that can resolve closely related bacteria within complex communities can lead to a better understanding of plant-microbe interactions. However, current methods lack the specificity to differentiate phylogenetically similar bacterial strains. In this study, we describe molecular strategies to study specific duckweed-bacteria interactions. We first systematically optimized a bead-beating protocol to co-isolate nucleic acids simultaneously from duckweed and bacteria. We then developed a generic fingerprinting assay to detect bacteria present in duckweed samples. To detect specific duckweed-bacteria interactions, we developed a genomics-based computational pipeline to generate bacterial strain-specific primers. These strain-specific primers differentiated bacterial strains from the same genus and enabled the detection of specific duckweed-bacteria interactions present in a community context. Moreover, we used these strain-specific primers to quantify the bacterial colonization of duckweed by normalization to a plant reference gene and revealed differences in colonization levels between strains from the same genus. Lastly, confocal microscopy of inoculated duckweed further supported our PCR results and showed bacterial colonization of the duckweed root-frond interface and root interior. The molecular methods introduced in this work should enable the tracking and quantification of specific plant-microbe interactions within plant-microbial communities.

## Introduction

The Lemnaceae, commonly known as duckweeds, is a family of freshwater aquatic plants [1]. Their small size, fast growth rate, growth habitat, and reduced morphology put forth duckweed as a model system to study plant-microbe interactions. Indeed, many similarities can be found between the structuring of duckweed-associated bacterial (DAB) communities and terrestrial plant bacterial communities. For example, both terrestrial plants and duckweed host distinct bacterial communities when compared to the host environment, demonstrating that selection strongly shapes the bacterial communities of both terrestrial plants and duckweed [2–5]. Moreover, similar bacterial taxa are found among terrestrial plant bacterial communities and DAB communities, suggesting bacterial adaptation to these plant habitats [3]. Therefore, studying duckweed-bacteria interactions may help reveal conserved mechanisms involved in plant-bacteria interactions.

The study of plant-bacteria interactions is complicated by many factors. One factor is the functional diversity found among phylogenetically similar bacteria associated with plants. Despite a similar phylogeny, these related bacteria can interact differently with plant hosts and may serve diverse roles in plant microbial communities. Community surveys of plant bacterial communities show that bacteria of the same genus can have different colonization dynamics across plant tissues and developmental stages [6]. Other community surveys show bacteria of the same family can have distinct plant host preferences [7]. In support of these community surveys, functional studies show bacterial strains from the same genus can colonize plants at different concentrations and protect against disease to different degrees [8,9]. Another factor that adds complexity to plant-microbe interactions is the presence of microbe-microbe interactions [10]. Bacteria may readily colonize plants when no other microbes are present. However, the same bacteria may not be able to stably colonize plants in a community context [11]. The presence of microbe-microbe interactions in microbial communities is a major reason why many bacteria display plant-growth-promoting behavior in the laboratory in mono-associations studies but not in field trials when natural microbial communities are present [12]. Thus, differentiating phylogenetically similar bacteria under diverse contexts will be important to unravel the complexity of plant-microbe interactions.

Current methods to study plant-bacteria interactions differ in the information they provide and in the context in which they can be applied [13]. The colony-forming units (CFU) assay is a classical microbiology technique used to quantify bacteria. In the context of plant-microbe interactions, this method has typically been used to quantify the colonization of plants by single bacterial isolates [14–17]. With the implementation of selective culture media, members in a small plant-bacterial community can also be distinguished [18]. However, this method can be laborious, imprecise, and cannot be used to quantify bacteria found in complex microbial communities. In contrast to the CFU assay, microscopy is a qualitative approach used to observe the spatial and temporal colonization dynamics of bacteria on plants [19]. Its application has revealed the presence of colonization hotspots on plants and different colonization patterns between bacteria when applied individually onto plant tissues [20–22]. However, microscopy commonly uses generic stains, fluorescent dyes, or oligonucleotide probes that cannot detect specific bacteria and may not be applicable for characterizing specific interactions within a bacterial community [23,24]. An alternative microscopy approach involves labeling and monitoring a bacterial strain of interest with an *in vivo* reporter gene, such as GFP or GUS, but this application can be laborious and is dependent on the transformability of the bacterium of interest [25–27]. Thus, the CFU assay and microscopy methods are often used to study bipartite plant-bacteria interactions, since they lack the specificity required to study the interactions between plants and specific members in complex bacterial communities. The most common method to detect bacteria in complex communities is 16S rRNA amplicon sequencing, in which variable regions of the 16S rRNA gene are selectively amplified and sequenced by high-throughput methods [28–30]. Initially, 16S rRNA amplicon sequencing provided the relative abundance of bacteria within communities but recent innovations allow for the absolute abundance of community members to be quantified [31–33]. Despite these improvements, this approach is still limited by the extent of polymorphisms in the 16S rRNA gene, which distinguishes between bacterial families and genera but lacks resolution between closely related bacterial species or strains of the same species [34–36]. In addition, some bacterial taxa contain multiple non-identical copies of the 16S rRNA gene, further complicating the differentiation of closely related bacteria with this approach [37,38]. As a result, no straightforward methods exist to study specific plant-bacteria interactions within complex communities.

To address this technical challenge in studying plant-bacteria interactions, we developed molecular methods to detect and characterize the colonization of duckweed by specific bacterial isolates under simple (i.e. binary) or community contexts. To apply molecular methods for the detection of duckweed-bacteria interactions, we first systematically optimized a bead-beating protocol to co-isolate nucleic acids simultaneously from both duckweed and bacteria. Second, we combined ribosomal intergenic spacer analysis (RISA) and PCR of a plant-specific marker to detect bacteria associated with duckweed. Third, we developed and implemented custom computational pipelines that can design primers to detect and quantify the colonization of duckweed by specific bacteria, either alone or in the presence of microbial communities. Lastly, we used confocal microscopy as a complementary approach to describe the bacterial colonization dynamics of *Lemna minor*. The molecular methods introduced in this work should enable high-resolution, quantitative studies of duckweed-bacteria interactions in diverse contexts.

## Results

### Selection of duckweed strain and bacteria isolates

Duckweed and bacteria were obtained to study duckweed-bacteria interactions. The duckweed strain, *Lemna minor* 5576 (Lm5576), was acquired from the Rutgers Duckweed Stock Cooperative (RDSC; New Brunswick, NJ, USA). This duckweed strain has been previously used to study duckweed-associated bacterial communities [3]. Bacteria originating from different hosts were acquired (File S1). One of the duckweed-associated bacterial (DAB) isolates, *Microbacterium sp*. RU370.1 (DAB 1A), was isolated from Lm5576 and can produce the phytohormone indole-3-acetic acid (IAA) as well as colonize and affect the root development of *Arabidopsis thaliana* [39,40] Another bacterial isolate was retrieved from the seaweed *Ulva fasciata*. This seaweed-associated bacterium, *Bacillus simplex* RUG2-6 (G2-6), was hypothesized to be a weak colonizer of duckweed due to the large evolutionary divergence between macroalgae and angiosperms. Two bacterial isolates of well-characterized plant colonizers, an epiphyte *Azospirillum brasilense* Sp7 (Sp7) and a known endophyte *Azospirillum baldaniorum* Sp245 (Sp245), were acquired to act as positive colonization controls [41]. Both these strains (Sp7 and Sp245) contain similar 16S rRNA gene sequences (97-99.9 % identity) and were initially classified as *Azospirillum brasilense*, but recent phylogenomic analyses show Sp245 belongs to the novel species named *Azospirillum baldaniorum* [42]. Together, these bacteria were used to inoculate duckweed to study their colonization dynamics of axenic Lm5576.

### Systematic optimization of a nucleic acid extraction method for duckweed-bacteria associations

To characterize bacterial colonization of duckweed using molecular methods, an optimized protocol for isolating nucleic acid simultaneously and efficiently from duckweed and different bacteria was developed. First, nucleic acid extraction was compared between bead-beating and homogenization with mortar and pestle using a modified CTAB protocol [43] (Figure S1). While mortar and pestle extracted more nucleic acids from Lm5576 than bead-beating, only bead-beating was able to extract both duckweed and bacterial nucleic acids. Therefore, bead-beating was selected as the physical lysis method for nucleic acid extraction. Various parameters of the bead-beating protocol were then modified to improve duckweed and bacteria nucleic acid extraction. First, three different sizes of beads were compared for their ability to extract plant and bacteria nucleic acid (Figure S2 and Figure S3). These tests showed 1.7 mm zirconium beads were the most effective in homogenizing duckweed tissues and extracting duckweed nucleic acids while 100 um silica beads were the best for extracting bacterial nucleic acids. Furthermore, a combination of different-sized beads effectively extracted nucleic acids from both duckweed and bacteria. Therefore, a combination of different-sized beads was used for the bead-beating protocol. Chloroform and a heating step at 65°C were then added to the lysis step to test their ability to improve nucleic acid extraction (Figure S4A). Both chloroform and heating improved nucleic acid extraction from duckweed. Bead-beating was also performed at different temperatures with and without the addition of a reducing agent to test for improvement of nucleic acid extraction (Figure S4B). All conditions resulted in high yields of intact nucleic acids, but bead-beating at room temperature without a reducing agent showed a slightly higher yield of nucleic acids and higher molecular weight DNA. Lastly, nucleic acid extractions were performed using different incubation times in the lysis buffer on different bacteria, including isolates from monoderm (Firmicutes and Actinobacteria) and diderm (Proteobacteria) bacterial phyla (Figure S5). Nucleic acids were extracted from both monoderm and diderm bacteria, with nucleic acid yield increasing with longer incubation times in the lysis buffer for some isolates. From these experiments, an optimized bead-beating protocol was developed to effectively extract nucleic acids from duckweed inoculated with different bacteria.

### Establishment of a PCR-based DNA fingerprinting assay for duckweed-bacteria interactions

rRNA intergenic spacer analysis (RISA) is a PCR-based method that amplifies the intergenic spacer region between 16S and 23S rRNA genes. This region can vary in copy number, sequence, and length between bacterial species. As a result, RISA can be used to estimate bacterial community composition by generating community fingerprints [44] and for rapid, universal bacterial typing [45]. In this study, RISA was applied as a simple molecular approach to detect the presence of different bacteria in association with duckweed. Different RISA primer sets were tested for their ability to amplify DNA from different bacterial species and duckweed (File S2, Figure S6). 16S-e1390f and 23S-e130r primers produced distinct fingerprints between bacterial species while they did not show amplification products from Lm5576 DNA under our conditions. Therefore, these primers were selected for detecting bacterial colonization of Lm5576. In addition to RISA, a plant-specific marker was used to compare the relative amount of Lm5576 genomic DNA between samples and to control for sample quality. Primers were designed for detecting the single-copy *Lemna minor* ortholog of the plant-specific *LEAFY* gene (*LmLFY*), which is a master transcription factor for flowering control (File S3). PCR using *LmLFY* primers specifically detected and allowed visual estimation of the relative quantity of Lm5576 DNA between samples (Figure S7). Both RISA PCR and *LmLFY* PCR were used in concert to monitor bacterial colonization of Lm5576. This strategy is subsequently referred to as “attachment PCR”.

### Standardization and validation of attachment PCR assay for molecular detection of duckweed-bacteria associations

Attachment PCR was used to detect and compare the colonization of Lm5576 by G2-6, DAB 1A, Sp7, and Sp245 bacterial strains (Figure 1A). Axenic Lm5576 plants were inoculated with bacteria for seven days. After seven days, inoculated Lm5576 tissue was collected, rinsed with sterile water, and nucleic acids were isolated using the bead-beating protocol described above. Isolated DNA from pure bacterial cultures and sterile Lm5576 were used as controls to compare RISA fingerprints from inoculated Lm5576 samples. RISA PCR did not generate any banding pattern from axenic Lm5576 DNA, whose sterility was verified by culturing on solid bacterial growth media (Materials and Methods). RISA PCR of bacteria DNA controls produced distinct fingerprints between G2-6, DAB 1A, and *Azospirillum* strains (Sp7 and Sp245). *LmLFY* PCR of DNA controls only produced a PCR product from the axenic Lm5576 DNA control and none from bacterial DNA controls. All inoculated Lm5576 samples showed *LmLFY* PCR products, indicating good sample quality and the presence of Lm5576 DNA for reference. RISA PCR of Lm5576 inoculated with G2-6 sample did not generate any bacterial fingerprint, suggesting G2-6 was either not able to colonize Lm5576 or colonized Lm5576 at a low concentration not detectable by RISA PCR. RISA PCR of Lm5576 inoculated with DAB 1A produced a fingerprint consisting of a single PCR band that matched the fingerprint of DAB 1A DNA, demonstrating DAB 1A colonized Lm5576. RISA PCR of Lm5576 inoculated with Sp7 or Sp245 produced a similar fingerprint consisting of two major PCR bands. These two bands were the most prominent PCR bands found in RISA PCR of Sp7 and Sp245 DNA controls indicating Sp7 and Sp245 were both able to colonize Lm5576. The higher concentrations of DNA used for Sp7 and Sp245 DNA controls may explain why the additional PCR bands were not clearly observed in Lm5576 inoculated with Sp7 or Sp245. Overall, attachment PCR showed that DAB 1A, Sp7, and Sp245 colonized Lm5576 at detectable levels. The exact matching of RISA PCR fingerprints between inoculated Lm5576 samples and DNA controls confirmed the colonization of Lm5576 by the respective bacteria. In addition, this exact matching suggested no contaminating bacteria were present. Therefore, fingerprint matching between RISA PCR of inoculated duckweed and DNA controls can be used to confirm what bacteria are present in duckweed samples.

**Figure 1.**
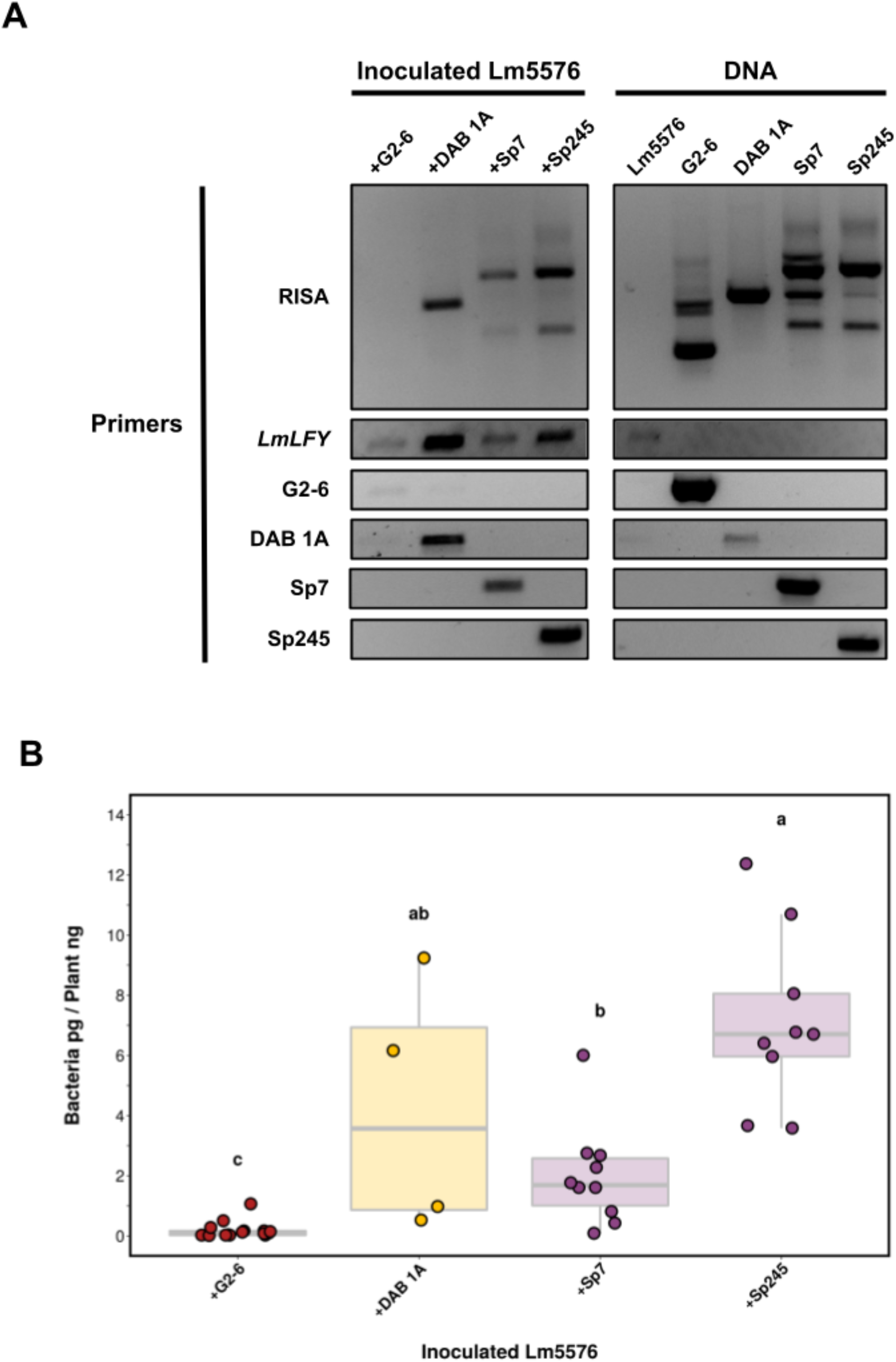
Molecular detection and quantification of specific duckweed-bacteria interactions. Lm5576 was inoculated with different bacteria in 0.5X SH. After seven days, inoculated Lm5576 tissue was collected, washed with sterile water, and nucleic acid was isolated for analysis. **A)** Representative gel electrophoresis results of end-point PCR using RISA, *LmLFY*, and strain-specific primers (File S2). RISA PCR fingerprints from inoculated Lm5576 samples were compared to the respective DNA controls from Lm5576 and bacteria alone. +G2-6 = Lm5576 inoculated with *Bacillus simplex* RUG2-6; +DAB 1A = Lm5576 inoculated with *Microbacterium sp*. RU370.1; +Sp7 = Lm5576 inoculated with *Azospirillum brasilense* Sp7; +Sp245 = Lm5576 inoculated with *Azospirillum baldaniorum* Sp245; RISA = PCR using 16S-1390f and 23S-e130r primers; *LmLFY* = PCR using LmLFY-F and LmLFY-R primers specific to Lm5576; DAB 1A = PCR using AmRU370.1-F and AmRU370.1-R primers specific to *Microbacterium sp*. RU370.1; G2-6 = PCR using BsRUG2.6-F and BsRUG2.6-R primers specific to *Bacillus simplex* RUG2-6; Sp7 = PCR using AbSp7-F and AbSp7-R primers specific to *Azospirillum brasilense* Sp7; Sp245 = PCR using AbSp245-F and AbSp245-R primers specific to *Azospirillum baldaniorum* Sp245. **B)** Bacterial colonization of Lm5576 was quantified using real-time PCR. Bacterial load was determined for each inoculated Lm5576 sample (picograms) and normalized to the amount of Lm5576 DNA in each sample (nanograms). Different colors were used for the different bacterial genera. Each data point represents an experimental repeat except for +G2-6, where each sample was measured twice. A significant difference was found in colonization loads between bacteria (Kruskal-Wallis, p-value = 4.28×10^−6^). Pairwise comparisons were performed using Dunn’s test and displayed as compact letters. Bacteria with significantly different colonization levels from each other, according to Dunn’s test, do not share any letters.

### Computational pipeline for primer design to detect and quantify specific duckweed-bacteria associations

Attachment PCR using RISA and *LmLFY* primer sets detected the colonization of Lm5576 by different bacteria, but it was unable to differentiate strains of the same genus (*i*.*e*., Sp7 and Sp245). To distinguish between closely related bacteria, a genomics-enabled approach was taken where strain-specific primers for traditional PCR were designed for each bacterium using available computational pipelines (File S2). For this approach, genomes of G2-6 and DAB 1A were sequenced and the genomes of Sp7 and Sp245 were retrieved from public databases (File S1). The strain-specific primers designed from this pipeline were used for PCR of DNA controls to validate their specificity (Figure 1A). As expected, strain-specific PCR of DNA controls uniquely detected the target bacteria and differentiated Sp7 and Sp245 strains. Strain-specific PCR of inoculated Lm5576 samples showed DAB 1A, Sp7, and Sp245 significantly colonized Lm5576. G2-6 specific PCR showed a faint amplification product in the Lm5576 sample inoculated with G2-6, in contrast to RISA PCR results, suggesting G2-6 attached to Lm5576 tissues at a low concentration that was not detectable by RISA PCR. These results show that PCR using bacterial strain-specific primers can uniquely detect phylogenetically similar bacterial strains and can be used to detect specific duckweed-bacteria interactions.

While end-point PCR with strain-specific primers detected specific duckweed-bacteria interactions, it could not be used to accurately quantify average bacterial colonization levels. To quantify bacterial colonization of Lm5576, bacterial strain-specific primers and Lm5576-specific primers were designed for real-time PCR (qPCR) assays using a custom computational pipeline (Methods, Figure S8, File S4). For this computational pipeline, unique genomic sequences were identified and retrieved for each bacterial genome. These unique sequences were then used for optimal primer design. qPCR with bacterial strain-specific primers from this pipeline was used to determine bacterial load for each inoculated Lm5576 sample. Bacterial abundance was then normalized to the quantity of Lm5576 DNA, which was determined using Lm5576-specific primers, for each inoculated Lm5576 sample. This approach is referred to as “attachment qPCR”. Attachment qPCR showed a significant difference (Kruskal-Wallis, p-value = 4.28×10^−6^) in the colonization of Lm5576 between the bacteria tested (Figure 1B). Attachment qPCR showed G2-6 colonized Lm5576 in significantly lower concentrations compared to the other bacteria tested (Dunn’s test, p-value < 0.005 for all comparisons), similar to what was found qualitatively by end-point strain-specific PCR (Figure 1A). DAB 1A and Sp245 had the highest bacterial colonization loads of Lm5576. However, DAB 1A displayed high variability between samples so no significant difference was established compared to Sp7 and Sp245. Sp7 colonized Lm5576 at significantly lower concentrations than Sp245 (Dunn’s test, p-value < 0.05). In conclusion, attachment qPCR revealed a significant difference in colonization levels between bacterial isolates from plants compared to the bacterial isolate from seaweed and detected significant differences in colonization levels between phylogenetically similar bacteria. These findings demonstrate attachment qPCR can be used to quantify colonization levels of bacteria on plants with high resolution.

### Bacterial colonization of Lemna minor visualized by confocal microscopy

As a complementary approach to the PCR-based approaches described above, confocal microscopy was performed on inoculated Lm5576 samples to qualitatively describe bacterial colonization patterns (Figure 2, File S5). Attachment PCR was performed on all microscopy samples and confirmed the colonization of Lm5576 by the respective bacteria and the absence of contaminating bacteria (File S6). All the bacteria tested were found to colonize the surface of Lm5576 fronds (Figure 2). G2-6, DAB 1A, and Sp245 were spread over the surface of Lm5576 fronds in smaller colonies while Sp7 was mostly localized to the root-frond interface in aggregates. No bacteria were observed to colonize the inside of Lm5576 fronds in these experiments. Bacteria displayed different colonization patterns of Lm5576 roots (Figure 2, File S5). G2-6 was found throughout the surface of Lm5576 roots at a low density. As mentioned above, Sp7 aggregates were mostly located on the surface of Lm5576 roots near the root-frond interface. DAB 1A and Sp245 were also found mostly at the root-frond interface on the surface of Lm5576 roots. Microscopy showed DAB 1A was present in higher concentrations at the root-frond interface than the other bacteria tested. This correlates with the high bacterial colonization load observed in attachment qPCR experiments for some samples (Figure 1B). Interestingly, Sp245 was found inside Lm5576 roots, within the endodermis, at high concentrations. This also agreed with the attachment qPCR results that revealed a significantly high colonization load for Sp245 and shows it is an endophyte for Lm5576. DAB 1A and Sp7 were also sporadically found inside Lm5576 roots but at a much lower frequency and concentration. In summary, confocal microscopy of inoculated Lm5576 samples revealed that the bacteria tested were able to colonize Lm5576 fronds and roots to various extents, with the root-frond interface as a hotspot for bacterial colonization. Of the 4 bacteria examined with confocal microscopy, G2-6 displayed the least amount of attached bacteria to Lm5576 while Sp245 showed the highest level of colonization, especially in the endosphere of the roots. These results support the main conclusions of the attachment PCR experiments in this study (Figure 1B) and contribute to the understanding of spatial colonization dynamics of bacteria on duckweed.

**Figure 2.**
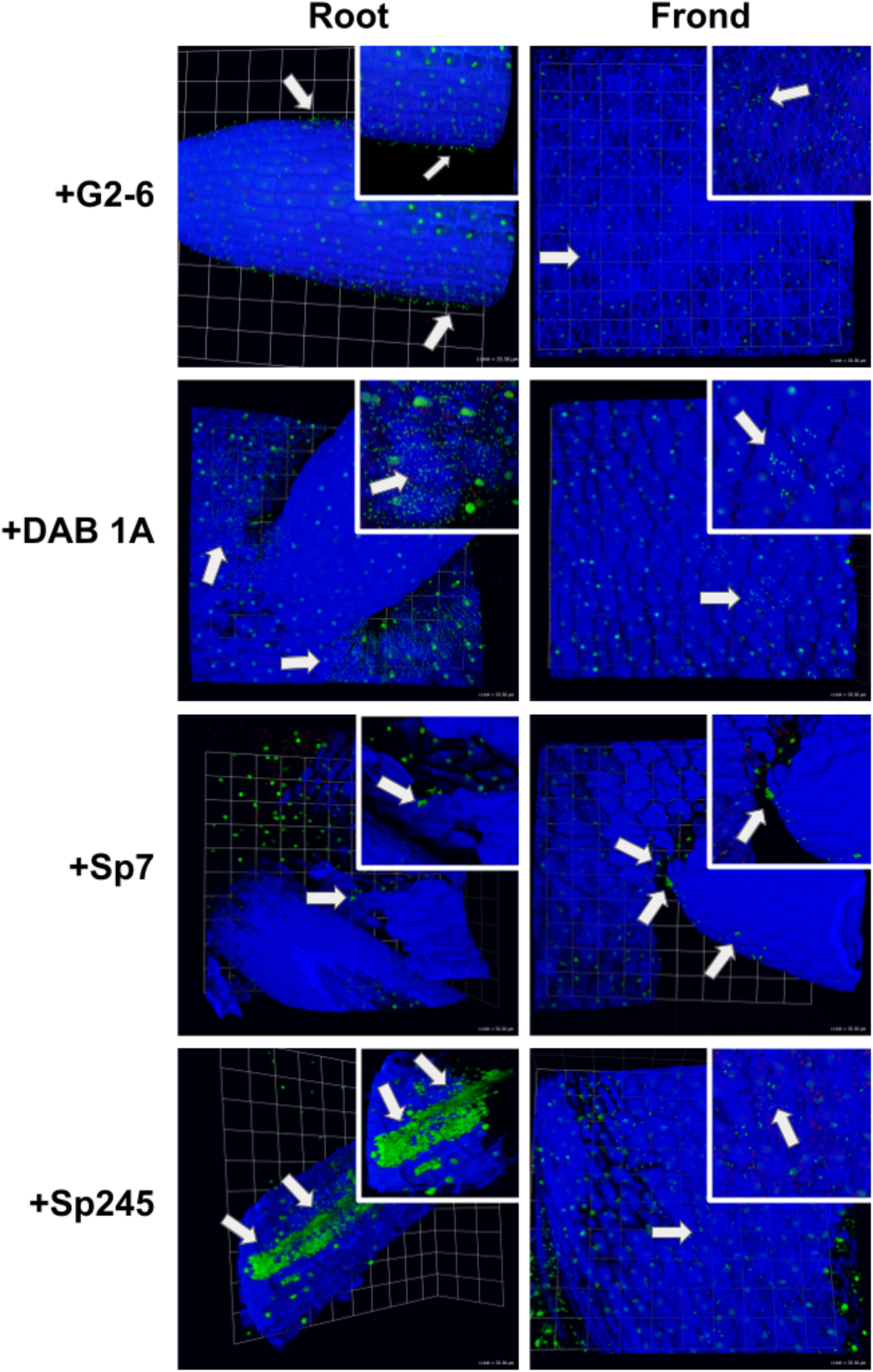
2D Confocal microscopy of inoculated duckweed samples. Confocal microscopy (40X/1.1 objective) was performed on inoculated Lm5576 in 0.5X SH media to spatially characterize bacterial colonization dynamics of duckweed. Calcofluor white was used to stain plant cellulose and visualized with the blue channel, SYBR Gold was used to stain DNA and visualized with the green channel, and chlorophyll autofluorescence was visualized with the red channel. Bacterial cells are stained green and are smaller in size compared to plant nuclei. For each image, white arrows point to cells of the respective bacterium, scale units are depicted in the bottom-right corner, and zoomed-in images are pictured in the top-right corner. +G2-6 = Lm5576 inoculated with *Bacillus simplex* RUG2-6; +DAB 1A = Lm5576 inoculated with *Microbacterium sp*. RU370.1; +Sp7 = Lm5576 inoculated with *Azospirillum brasilense* Sp7; +Sp245 = Lm5576 inoculated with *Azospirillum baldaniorum* Sp245.

### Strain-specific monitoring of duckweed-bacteria associations in a community context

To further illustrate the efficacy of attachment PCR, this method was used to detect specific duckweed-bacteria interactions in the presence of other bacterial isolates and in the presence of microbes in wastewater (Figure 3). Attachment PCR was also tested using another duckweed strain obtained from the RDSC, *Spirodela polyrhiza* strain 9509 (dw9509), whose genome has been sequenced to reference quality [46,47]. For these experiments, dw9509 was inoculated with DAB 1A, *Bacillus sp*. RU9509-4 (DAB 3D), Sp245, and wastewater containing microbes for five days. In addition, Sp245 was co-inoculated onto dw9509 with either DAB 1A, DAB 3D, or wastewater containing microbes to test bacterial colonization in the presence of other microbes. After five days, inoculated duckweed tissue was collected, rinsed with sterile water, and nucleic acids were isolated. *SpLFY* PCR generated a PCR product for all inoculated dw9509 samples, ensuring good sample quality. RISA PCR and strain-specific PCR did not generate any signals for axenic dw9509, confirming its sterility. RISA PCR showed DAB 1A, DAB 3D, Sp245, and wastewater microbes colonized dw9509. Additionally, strain-specific PCR confirmed DAB 1A, DAB 3D, and Sp245 colonized dw9509, while no amplification product was obtained with wastewater containing microbes. RISA PCR and Sp245 strain-specific PCR demonstrated Sp245 was able to colonize dw9509 in the presence of DAB 1A, DAB 3D, and a wastewater microbial community, indicating robust colonization ability by Sp245 under diverse contexts. While both DAB 1A and DAB 3D were able to colonize dw9509 in the presence of Sp245, DAB 3D strain-specific PCR showed a lower amplification signal in the dw9509 sample co-inoculated with DAB3D and Sp245 compared to the dw9509 sample inoculated only with DAB 3D, suggesting DAB 3D colonization of dw9509 was reduced in the presence of Sp245. These experiments illustrate the efficacy of attachment PCR and strain-specific PCR to detect specific duckweed-bacteria interactions in a community context. In addition, quantitative effects on bacterial colonization of the plant host resulting from microbe-microbe interactions can be revealed.

**Figure 3.**
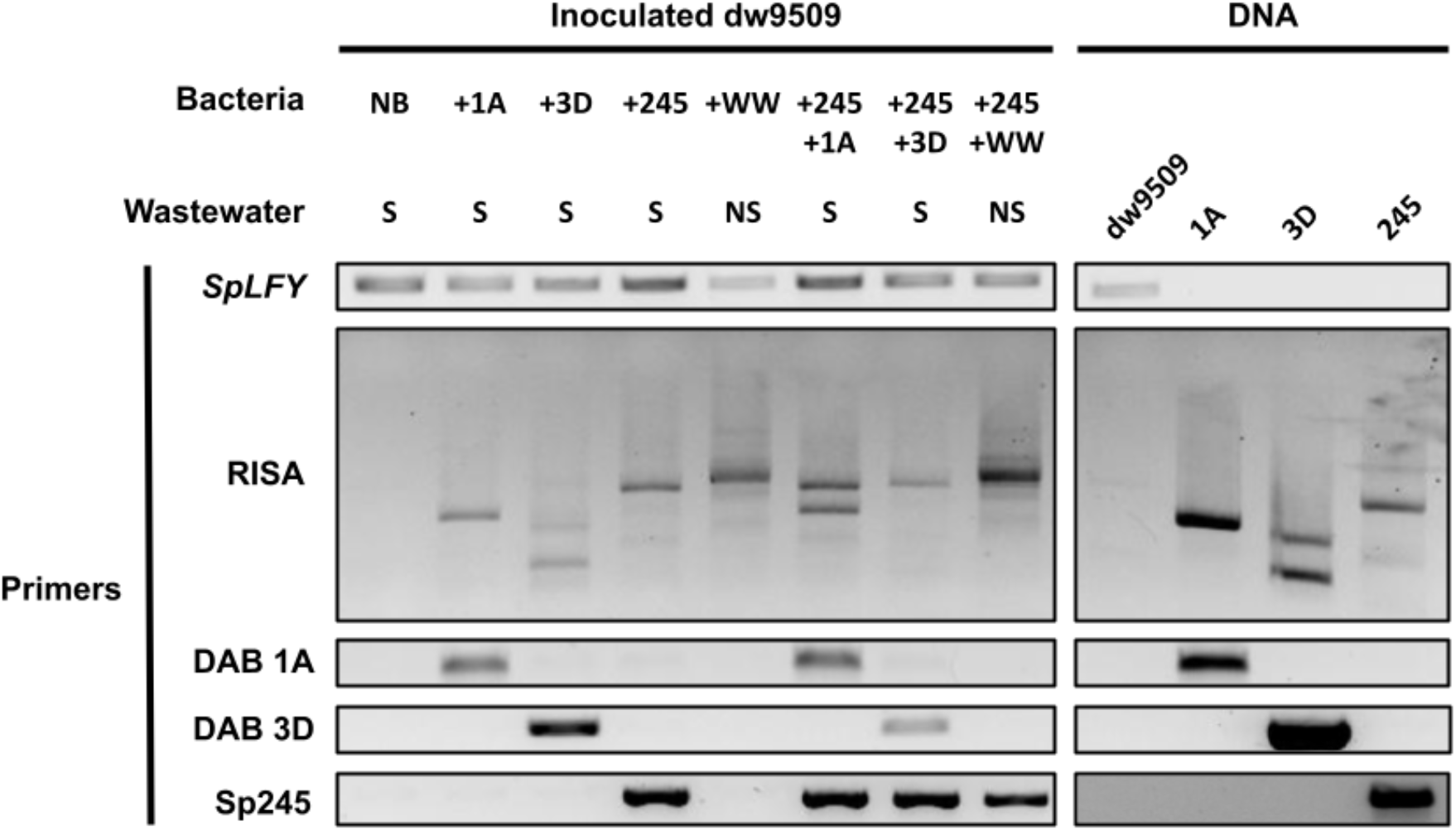
Molecular detection of specific duckweed-bacteria interactions in a community context. dw9509 was inoculated with different bacteria in wastewater with or without microbes. In addition, Sp245 was co-inoculated onto dw9509 with DAB isolates or non-sterile wastewater containing microbes. For co-inoculated samples, bacteria were mixed at a 1:1 ratio. After five days, dw9509 tissue was collected, washed with sterile water, and nucleic acids were isolated for end-point PCR using RISA, *SpLFY*, and strain-specific primers (File S2). RISA PCR fingerprints from dw9509-bacteria samples were compared to DNA controls from dw9509 and bacteria alone. Wastewater: S = filter-sterilized wastewater not containing microbes, NS = non-sterile wastewater containing microbes; Bacteria: NB = axenic dw9509, +1A = dw9509 inoculated with *Microbacterium sp*. RU370.1, +3D = dw9509 inoculated with *Bacillus sp*. RU9509.4, +Sp245 = dw9509 inoculated with *Azospirillum baldaniorum* Sp245, +WW = dw9509 inoculated with non-sterile wastewater containing microbes; Primers: *SpLFY* = PCR using SpLFY-F and SpLFY-R primers specific to dw9509, RISA = PCR using 16S-1390f and 23S-e130r primers, DAB 1A = PCR using AmRU370.1-F and AmRU370.1-R primers specific to *Microbacterium sp*. RU370.1, DAB 3D = PCR using BsRU9509.4-F and BsRU9509.4-R primers specific to *Bacillus sp*. RU9509.4, Sp245 = PCR using AbSp245-F and AbSp245-R primers specific to *Azospirillum baldaniorum* Sp245

## Discussion

### Localization of bacteria on duckweed

In terrestrial plants, bacteria have been shown to consistently colonize certain areas of plants termed colonization hot spots [20], which include root cracks where lateral roots emerge from the main root [21]. One explanation for this bacterial colonization pattern is that root cracks may release cell lysates and exudates that could help attract bacteria and other microorganisms [48]. In duckweed, a few studies have already described bacterial colonization patterns of duckweed. In one study, duckweed collected from chalk streams was found to have a higher density of bacteria on the submerged abaxial surface of duckweed fronds compared to the aerial adaxial surface [49]. In another study on *L. minor*, the plant-growth-promoting bacterium, *Bacillus amyloliquefaciens* FZB42, was found to initially colonize *L. minor* at the root tip and root-frond interface and later the grooves between root epidermal cells and concavities of the abaxial frond surface [27]. While for a rootless duckweed such as *Wolffia australiana*, bacteria present in the surrounding greenhouse environment were found to colonize *W. australiana* near reproductive pockets, where mother and daughter fronds attach, and the stomata [4,21]. In the present study, we performed high-resolution confocal microscopy on inoculated *L. minor* samples to further study bacterial colonization patterns of duckweed (Figure 2). All bacteria in this study were able to colonize the abaxial surface of duckweed fronds and roots to varying extents. Like previous reports [27], bacterial strains used in this study also showed a preference for the root-frond interface. Together these studies show that bacteria readily colonize the abaxial surface of duckweed fronds, at least for duckweeds with roots. One possible explanation for this observation is that the abaxial side of duckweed fronds is in direct contact with the microbial inoculum present in the surrounding water environment. Another explanation is surface composition, such as the cuticle, is distinct between the abaxial and adaxial surface of fronds [50] and may play a role in the differential attachment of microbes. Furthermore, while duckweeds do not make lateral roots [51], the root-frond interface in duckweed may serve as a hotspot akin to the root cracks in terrestrial plants, where cell contents are released or secreted to attract microbes.

While the surface colonization patterns of bacteria on duckweed have been described, there is no description of whether or not bacteria can colonize the inside of duckweed tissues. In this study, Sp245 colonized Lm5576 at the root-frond interface and the inside of duckweed roots, within the endodermis [51], at high densities (Figure 2). To the best of our knowledge, this is the first report of endophytic colonization in duckweed. In terrestrial plants, colonization hot spots, such as root cracks, can be used by bacteria to enter the roots of terrestrial plants [21,52,53]. Likewise in duckweed, one possibility could be that Sp245 entered through cracks at the root-frond interface of Lm5576 and proceeded to colonize the interior of Lm5576 roots. Recently, we studied the Sp245 interaction with the model plant *Arabidopsis thaliana* and revealed its potential interaction with guard cells in leaf tissues as a means for entering the endosphere [40]. Strikingly, this interaction and targeting to the guard cells by Sp245 is abolished in the pleiotropic *axr1* mutant, suggesting specific involvement of this gene in the signaling between plants and certain microbes. However, our microscopy studies with Lm5576 failed to observe this guard-cell colonization of Sp245 on duckweed fronds, indicating this mode of interaction could be lost or modified in duckweed. Sp245 was originally isolated from wheat experiments in Brazil [54] and colonizes the inside of wheat roots at high densities [22,24]. In contrast to the endophytic colonization pattern of Sp245, Sp7, originally isolated from *Digitaria* in Brazil [55], is an epiphyte shown to colonize only the surface of plant roots like wheat [22,24]. Sp7 aggregates were also found on the surface of corn roots under high culture concentrations [56]. Likewise, we found Sp7 mostly colonized the surface of Lm5576 roots near the root-frond interface in aggregates. These Sp7 and Sp245 colonization experiments on Lm5576 demonstrate that plant-associated bacteria can have similar colonization patterns with both terrestrial plants, like wheat, and aquatic monocots, like duckweed. This suggests a likely conservation of bacterial mechanisms for association with duckweed and other higher plants.

### Methods for molecular detection and quantification of specific duckweed-bacteria interactions

To date, studies of duckweed-bacteria interactions have relied on classical microbiology techniques like the CFU assay to monitor bacterial colonization of duckweed [57,58]. As mentioned above, this CFU assay lacks the specificity to differentiate bacteria within a community context. To enable the detection of specific duckweed-bacteria interactions, we decided to apply PCR-based approaches to characterize duckweed-bacteria interactions. A prerequisite for using such approaches is a protocol capable of efficiently isolating nucleic acids. Thus far, there have been no attempts to systematically develop a protocol capable of efficiently isolating nucleic acids from both duckweed and bacteria. A working protocol for isolating nucleic acids is critical for studying duckweed-bacteria interactions since different DNA extraction protocols can introduce significant biases toward what bacteria are detected and in what quantities [59,60]. These differences can be partly explained by the inability of certain protocols to efficiently lyse monoderm bacteria, gram-positive bacteria consisting of a thick peptidoglycan layer. However, nucleic acid isolation protocols implementing physical lysis methods such as bead-beating can efficiently lyse monoderm bacteria especially when longer bead-beating times are used [61][61–63][61]. Bead-beating protocols are also reproducible [60,64], yield high concentrations of nucleic acid [60,65], and produce more accurate community profiles [66]. In addition, combining bead-beating and chemical lysis, such as phenol or chloroform, can dramatically increase DNA extraction efficiency and quality [60,65]. For these reasons, bead-beating is recommended for nucleic acid extraction protocols [67]. Here, we implemented and optimized a bead-beating protocol to simultaneously co-isolate duckweed and bacteria nucleic acids. By combining different bead sizes and a CTAB/chloroform lysis buffer, this bead-beating protocol produced high yields of intact nucleic acids from duckweed and different bacteria, including monoderm and diderm bacteria (Figures S2-S5). Furthermore, through various testing of the bead-beating protocol, we observed increases in nucleic acid yields with longer incubation time periods in the lysis buffer (Figure S5) and with longer alcohol precipitation time periods. However, the ability of this bead-beating protocol to generate representative profiles of duckweed colonized by complex bacterial communities remains to be validated. This could be tested by isolating nucleic acid from mixtures of bacteria in known concentrations, known as mock communities [68,69]. This will be an important validation step for applying this bead-beating protocol to study the interactions between duckweed and complex microbial communities in the future. Lastly, this bead-beating protocol can be modified to isolate only DNA or RNA from duckweed or bacteria by adding an RNase or DNase treatment step respectively.

rRNA intergenic spacer analysis (RISA) has been commonly used for community fingerprinting [44] and bacterial typing [45]. However, RISA has also been used to study plant-bacteria interactions. For example, an automated version of RISA (ARISA), that applies fluorescently tagged primers and detects fluorescent PCR fragments [70], was used to monitor changes in the composition of synthetic bacterial communities [71]. Here we used RISA to detect bacterial colonization of duckweed by comparing fingerprints of inoculated duckweed samples to DNA controls of duckweed and the respective bacteria alone (Figure 1A, Figure 3). This molecular approach serves many purposes. First, RISA can be used to determine the axenic status of *L. minor* and *S. polyrhiza* plants used in experiments since RISA PCR does not produce any amplicons from sterile Lm5576 and dw9509. This is worth highlighting since difficulties can be encountered in obtaining sterile duckweed [72]. As we optimized RISA PCR for use with *L. minor* and *S. polyrhiza* in this study, RISA PCR may need to be optimized for use with other duckweed species, by using different RISA primer sets and/or PCR conditions. Second, RISA can be used to determine the colonization of duckweed by different bacterial species since RISA PCR can generate distinct fingerprints between bacterial species. Third, because fingerprints generated from inoculated duckweed samples are compared to DNA controls of the organisms being studied, RISA can also reveal non-matching fingerprints that are due to contaminating or exogenous bacteria. In addition to RISA, we included a duckweed*-*specific marker to control for sample quality as well as to provide a reference for the relative quantity of duckweed DNA between samples. We termed this approach, of combining RISA PCR and PCR of a plant-specific marker, as attachment PCR. Attachment PCR was recently used in our lab to detect interactions between bacteria isolated from rice and duckweed [73]. Attachment PCR under laboratory conditions showed that *Pantoea* isolates from rice were able to colonize duckweed such as Lm5576, despite the low representation of *Pantoea* bacteria in duckweed-associated bacterial communities from the same rice paddies. This suggested that microbe-microbe interactions or environmental factors could be responsible for the low representation of *Pantoea* in duckweed-associated bacterial communities in this context. These case studies demonstrate the utility of RISA in general and its application for attachment PCR specifically. Attachment PCR, as a molecular approach, is a more quantitative and specific method than microscopy or bacteria counting and could lead to more mechanistic analyses of plant-microbe interactions as we have shown before [73].

Despite its advantages, we found RISA was unable to distinguish between strains from the same genus (Figure 1A). As mentioned previously, plant-associated bacteria from the same genus can be functionally diverse. Thus, it is important to identify methods that can differentiate closely related bacteria isolated from plants. Here, we used available computational tools as well as developed a custom computational pipeline to generate strain-specific primers, leveraging the large and growing databases for bacteria (Figure S8, File S2, File S4). These strain-specific primers were able to clearly differentiate strains from the same genus (Figure 1A). In combination with the attachment PCR approach, strain-specific PCR can provide a more complete description of the duckweed-bacteria interactions present in samples. One application is that this combinatorial approach can be used to ensure reproducible duckweed-bacteria interactions between experiments.

Strain-specific primers can also be designed for real-time PCR to quantify specific bacterial colonization of Lm5576 by normalizing to a duckweed-specific reference marker (Figure 1B). This “attachment qPCR” approach, of normalizing bacterial colonization load to an internal plant marker gene, has been applied in previous studies to quantify plant root colonization by arbuscular mycorrhizal fungi [74], Rhizobiales re-colonization of plant roots [75], and bacterial abundance in the phyllosphere of *Arabidopsis thaliana [71]*. However, these studies used approaches that catered to the specific purposes of these experiments or did not provide an accessible strategy to design primers. Here, we developed a straightforward computational pipeline to design strain-specific primers for any bacterial strain with a sequenced genome. While this computational pipeline was used in this study to design primers to characterize bacterial colonization of duckweed, the pipeline could be readily applied to other host systems.

### Bacterial adaptation to plant habitats and colonization dynamics

Selection is a major driver in structuring plant bacterial communities [2]. As a result of this selection, certain bacteria have adapted to occupy different plant habitats [76,77]. For example, genomic analyses have shown that plant-associated bacterial genomes are enriched in certain functions like chemotaxis, motility, and carbohydrate metabolism [78,79]. In support of these analyses, genome-wide functional screens, using transposon sequencing, in both terrestrial plants and duckweed confirm the involvement of chemotaxis, motility, and carbon metabolism in bacterial colonization of plants [80,81]. In addition to these functions, many plant-associated bacteria are capable of producing phytohormones, such as auxins, which can have either beneficial or detrimental effects on plant hosts [82]. Most studies on bacterial auxin production have focused on the effects on plant growth, but one recent study investigated the role of bacterial auxin production in plant colonization [83]. This study showed that bacterial auxin production is necessary for efficient root colonization for some bacteria and revealed a feedback loop between auxin-producing bacteria and the plant host. In this feedback loop, auxin-producing bacteria elicit an immune response from the plant host that produces reactive oxygen species (ROS). These ROS induce auxin production in bacteria, where the bacterial auxin detoxifies the ROS from the plant host. This ROS detoxification allows bacteria to efficiently adhere and form colonies on plant roots. In turn, this bacterial colonization further elicits ROS production by the plant host immune response. Together, these studies describe some of the functions that have evolved in bacteria to colonize plants.

In this study, we explored the colonization levels among a non-plant-associated bacterial isolate and several plant-associated bacterial isolates using attachment qPCR. G2-6 was isolated from seaweed, a macroalga from salt water, and likely has not adapted or evolved to colonize freshwater macrophytes like duckweed. We thus expected G2-6 to colonize duckweed at very low levels, if at all. Indeed, attachment qPCR showed G2-6 colonized Lm5576 at significantly lower concentrations compared to all the plant-associated bacterial isolates tested (Figure 1B). DAB 1A was originally isolated from Lm5576 and produces high levels of the auxin indole-3-acetic acid (IAA) that affects the root development of *Arabidopsis thaliana* [39,40]. Therefore, we expected DAB 1A to re-colonize Lm5576 in this study. Confocal microscopy confirmed these expectations and showed high levels of DAB 1A near the root-frond interface of Lm5576 (Figure 2). While attachment qPCR showed variable colonization levels of DAB 1A, DAB 1A colonized Lm5576 at high levels in some samples (Figure 1B). Extending these results, confocal microscopy of *A. thaliana* inoculated with DAB 1A showed high concentrations of DAB 1A present on the root surface [40]. Interestingly, this same study showed another DAB isolate, DAB 33B, was not able to colonize *A. thaliana* even though it belonged to the same genus, *Microbacterium*, as DAB 1A. In addition to this inability to colonize *A. thaliana*, DAB 33B was shown to produce significantly lower levels of IAA compared to DAB 1A. Together, one explanation for the different colonization dynamics between these phylogenetically similar strains (DAB 1A and DAB 33B), could be that high levels of auxin production facilitate DAB 1A colonization of plants. Members of the *Azospirillum* genus are well-known plant colonizers and have been shown to fix nitrogen and produce phytohormones, such as IAA, that may promote plant growth [41]. Interestingly, *Azospirillum* taxa have also been detected in surveys of DAB communities [3] as well isolated from duckweed tissues [84]. Therefore, we hypothesized Sp7 and Sp245, both *Azospirillum*, would be able to colonize Lm5576 to some extent. Attachment qPCR showed Sp245 colonized Lm5576 at significantly higher levels than Sp7 (Figure 1B). This was further supported by confocal microscopy which showed significantly high concentrations of Sp245 within duckweed roots (Figure 2). These results are similar to a previous study that found Sp245 colonized the root endosphere of wheat and contained higher overall colonization levels compared to Sp7 [22].

Together, these attachment qPCR results raise several implications about the bacterial colonization of plants. For one, these data suggest that bacteria adapted to plants may display significantly higher colonization levels compared to non-adapted bacteria (Figure 1B). If so, then attachment qPCR can be used to screen for bacteria adapted to colonize plant habitats. This kind of experiment may help to discover novel traits necessary for the successful bacterial colonization of plants. Secondly, the plant-associated bacterial isolates examined in the present work showed different colonization levels. This raises the question, what traits determine the colonization levels of bacteria on plants? As mentioned above, auxin production is necessary for some bacteria to colonize plants [83]. Interestingly, the plant-associated bacterial isolates DAB 1A, Sp7, and Sp245 all produce significant levels of auxin [39,40]. Future work could use attachment qPCR to examine the relationship between the levels of bacterial auxin produced and the effect on bacterial colonization levels of plants. Results from this study also showed significantly higher colonization levels for the endophyte Sp245 compared to the epiphyte Sp7 (Figure 1B). This also raises the question, what is the relationship between bacterial colonization levels and bacterial colonization patterns? Do all endophytes display high colonization levels? If not, what controls the colonization levels of different endophytes? To answer this, attachment qPCR experiments could be performed to quantify the colonization levels on different bacterial endophytes. In summary, quantitative studies using attachment qPCR could lead to an improved understanding of traits underlying the bacterial colonization of plants.

### Detection of specific duckweed-bacteria interaction within a community context

Similar to findings with terrestrial plant bacterial communities, microbe-microbe interactions likely play a role in bacterial colonization of duckweed. One study reported that a plant-growth-promoting bacterium (PGPB) and two different plant-growth-inhibiting bacteria (PGIB) showed stable colonization levels of duckweed when inoculated separately [58]. However, when the PGPB and PGIB were co-inoculated together, the PGPB strain completely excluded one of the PGIB from colonizing duckweed. In another study, the same PGPB strain slowly decreased in abundance over time on duckweed in the presence of different bacterial communities [85]. Thus, the ability to distinguish between phylogenetically similar microbes in both mono-associations and within a community context will be important for studying the interactions between plants and complex microbial communities. In our work, strain-specific primers were shown to detect specific duckweed-bacteria interactions within a community context (Figure 3). The specificity demonstrated by strain-specific PCR has a pertinent application in the synthetic ecology approach used to study plant-microbe interactions [86]. In this approach, synthetic bacterial communities (SynComs) are constructed from bacterial isolates that are representative of members found in wild plant bacterial communities. In contrast to wild bacterial communities, SynComs are experimentally amenable and tractable allowing causal relationships to be determined. As a constructed community, SynComs can capture the complexity of plant bacterial communities found in nature while providing a means to decipher mechanisms underlying community dynamics and functions [19]. However, SynComs are limited by methods commonly used to track member presence and abundance, such as 16S rRNA amplicon sequencing. Since 16S rRNA amplicon sequencing can’t distinguish between many phylogenetically similar bacteria, SynComs have to be carefully designed in a way to select distinguishable members [87]. As a result, this can severely limit the diversity and representativeness of SynComs that can be used to effectively study the colonization dynamics of plant microbial communities. Using strain-specific primers will allow closely related bacteria to be included and monitored in SynCom experiments. Moreover, attachment qPCR can be used to quantify member abundance in SynCom experiments. The strategy used in attachment qPCR, where bacteria load is normalized to the quantity of host DNA, is similar to traditional qPCR used in RNA-sequencing experiments to validate gene expression, where a target gene is normalized to a housekeeping gene. In an analogous fashion, attachment qPCR could be used to compare member abundance generated from 16S rRNA amplicon sequencing in SynCom experiments since both approaches are DNA-based. Moreover, attachment qPCR could allow phylogenetically similar bacteria with different colonization dynamics and functional traits to be used in SynComs. Such comparisons could facilitate the assignment of the different phenotypes observed in SynCom experiments to specific molecular features. Together, these kinds of experiments should facilitate a mechanistic understanding of the interactions between plant hosts and their associated microbes.

## Conclusions

In conclusion, the PCR-based approaches introduced in this study have been shown to be effective for studying duckweed-bacteria interactions. Attachment PCR with generic RISA primers can be used to reveal the bacteria associated with duckweed while PCR using strain-specific primers can be used to differentiate specific duckweed-bacteria associations. Additionally, the attachment qPCR approach can be used to quantify colonization levels of bacteria under binary or community contexts. While these molecular approaches were used to study duckweed-bacteria interactions in this study, they should be easily adopted for use with other host-microbe systems. Together, these strain-specific approaches overcome the limitations of current methods used to detect plant-microbe interactions and enable the detection and quantification of specific plant-microbe interactions under diverse scenarios.

## Materials & Methods

### Duckweed sterilization and propagation

Cultures of *Lemna minor* 5576 (Lm5576) and *Spirodela polyrhiza* (dw9509) were obtained from the Rutgers Duckweed Stock Cooperative (RDSC; Rutgers University, New Brunswick, NJ, USA). Duckweed cultures were sterilized using a modified protocol from a previously described procedure [72]. For this procedure, duckweed plants were transferred to 1.7 mL microcentrifuge tubes and washed with 500 uL of salt and detergent solution (1 % Triton-X 100, 137 mM NaCl, 2.7 mM KCl, 10 mM Na2HPO, 1.8 mM KH2PO4, 0.5 mM MgSO4, 1 mM CaCl2, pH 7.4) to facilitate surface sterilization. Duckweed plants were then surface-sterilized using 5-10 % (v/v) household bleach (0.5-1 % sodium hypochlorite). Duckweed plants were surface sterilized until most frond tissues turned white and only the meristematic regions retained chlorophyll and remained green. Following surface sterilization, 2 % (w/v) of sodium thiosulfate was added to help neutralize residual sodium hypochlorite [88]. Surface-sterilized duckweed plants were then rinsed with sterile water and aseptically transferred to 0.8 % (w/v) agar (BD, Catalog #214530) plates with 0.5X Schenk and Hildebrandt basal salt mixture (SH) media (Phytotechnology Laboratories, Catalog #S816) containing 0.5 % sucrose and 100 ug/mL cefotaxime (GoldBio, Catalog #C-104-25) at pH 6.5-7.0. In addition, surface-sterilized duckweed plants were transferred to 1.5 % (w/v) agar plates with Miller’s (10 g/L tryptone, 5 g/L yeast extract, 10 g/L NaCl) lysogeny broth (LB). Surface-sterilized duckweed plants were allowed to propagate on the 0.5X SH agar plate and the LB agar plate. The LB agar plate was observed for any signs of microbial growth. If microbial growth was observed on duckweed plants growing on the LB agar plate then the surface-sterilization procedure was repeated on the surface-sterilized duckweed plants growing on the 0.5X SH agar plate.

Once axenic duckweed plants were obtained, stock cultures and working cultures of axenic duckweed were generated. Stock cultures of axenic duckweed were stored at 15°C and only used when required. Axenic working cultures of duckweed were generated by transferring a few duckweed plants to a 0.5X SH agar plate with 0.5 % (w/v) sucrose and an LB agar plate after each experiment. If no microbial growth was observed on the LB agar plate then duckweed plants on the 0.5X SH agar media were propagated for experiments. If microbial growth was observed, then a stock culture of axenic duckweed was retrieved from storage and propagated for experiments.

Axenic duckweed plants were propagated in a growth chamber on 0.5X SH agar media with 0.5 % (w/v) sucrose (pH 6.5-7.0) at 25°C under a photoperiod of 16 hours light and 8 hours dark for 2-4 weeks. Duckweed plants from the agar plate were then transferred to a 100 mL liquid culture of 0.5X SH with 0.1 % (w/v) sucrose and propagated for 1-2 weeks under the same growth chamber conditions. Axenic duckweed plants from these liquid cultures were then transferred for experiments. Duckweed sterility was confirmed between transfers by plating duckweed plants on LB agar plates and checking for microbial growth.

### Bacteria isolation and identification

To inoculate duckweed with bacteria for experiments, bacteria were isolated from different duckweed samples and the seaweed *Ulva fasciata* by washing tissues before homogenization and plating on LB agar or tryptic soy agar (TSA; BD, Catalog #236950) plates at 28°C for 2 to 3 days (File S1). For some bacterial isolates, plant host tissues were surface-sterilized, using the procedure described above, before isolation. Pure cultures for these bacterial isolates were generated by picking single colonies from LB agar or TSA plates and inoculating liquid LB or tryptic soy broth (TSB; Hardy Diagnostics, Catalog #C7141) for up to two days at 28°C. Glycerol stocks were then generated for each isolate and stored at -80°C as stock cultures until further use. Cultures of *Azospirillum* strains, *A. brasilense* Sp7 and *A. baldaniorum* Sp245 (formerly *A. brasilense*) [42], were obtained from S. Lebeis (MSU) and stored as glycerol stocks.

Bacterial isolates from duckweed and seaweed were previously identified using the following procedure [39]. The 16S rRNA gene fragment was amplified with polymerase chain reaction (PCR) using the primers 16S-e9f and 16S-e926r (File S2) [30]. PCR reactions were composed of 0.4 uM of each primer, 0.2 mM dNTPs, 2 units of Choice-Taq DNA polymerase in 1X NH4 reaction buffer (Thomas Scientific, Catalog #CB4050-2), and 1 uL of either bacterial nucleic acid (100 ng/uL), bacterial DNA (5 ng/uL), or bacterial liquid culture. PCR reactions were performed using the following 3-stage thermocycler program: 1) denaturation stage of 95°C for 5 minutes, 2) cycling stage of 25 cycles consisting of of 95°C for 30 seconds, 50°C for 30 seconds, 72°C for 1 minute, and 3) a final extension stage of 72°C for 5 minutes. PCR products were cleaned using ExoSAP-It PCR Product Cleanup Reagent (ThermoFisher Scientific, Catalog #78200.200.UL) or DNA Clean & Concentrator-5 kit (Zymo Research, Catalog #D4003). PCR products were sent to Genewiz (South Plainfield, NJ, USA) for sequencing using both 16S-e9f and 16S-e926r primers.

For each isolate, the resulting chromatograms for both forward and reverse sequences were analyzed and poor-quality sequences at both 5’ and 3’ ends were cropped using Geneious (www.geneious.com) or FinchTV v1.3.0 (Geospiza, Inc.)(www.digitalworldbiology.com). Forward and reverse sequences were aligned using SerialCloner v2.6.1 (http://serialbasics.free.fr/Serial_Cloner.html) to generate a consensus sequence. Gaps and mismatches were corrected in the consensus sequence using the chromatograms of the raw sequences. The consensus sequence was cropped 216 bp downstream and 385 bp upstream of the conserved U515 (5’-GTGCCAGCAGCCGCGGTAA-3’) sequence [30] to generate a 620 bp processed sequence. Processed sequences were annotated using the RDP classifier v2.13 with the 16S rRNA training set 18 [89].

### Bacteria genome sequencing

Draft genomes were generated at the DOE Joint Genome Institute (JGI) for duckweed-associated bacterial (DAB) isolates DAB 1A and DAB 3D as well the seaweed bacterial isolate G2-6 (File S1). Standard 300 bp Illumina shotgun libraries were constructed for all isolates.

DAB 1A (*Microbacterium sp*. RU370.1) and DAB 3D (*Bacillus sp*. RU9509.4) libraries were sequenced with the Illumina HiSeq 2000 platform. Raw reads were filtered for artifacts using BBDUK (Bushnell B., sourceforge.net/projects/bbmap/). Filtered reads were assembled using Velvet v1.2.07 [90] with the following parameters: velveth 63 -shortPaired, velvetg - very_clean yes -exportFiltered yes -min_contig_lgth 500 -scaffolding no -cov_cutoff 10. Velvet contigs were then used to create 1-3 kb simulated paired end reads using wgsim v0.3.0 (https://github.com/lh3/wgsim) with the following parameters: -e 0, -1 100, -2 100, -r 0, -R 0, -X 0. Simulated read pairs were then used to assemble Illumina reads using Allpaths-LG version r46642 [91] with the following parameters: PrepareAllpathsINputs, RunAllpathsLG. Assembly of 16S rRNA genes (percent 16S rRNA sequence covered in assembly is >= 80 % or length >= 1000 bp) was performed using filtered Illumina reads and non-duplicated sequences were merged into Allpaths assembly.

G2-6 (*Bacillus simplex* RUG2-6) libraries were sequenced with the Illumina HiSeq-2500 1TB platform. Read were processed using the BBTools suite at JGI (BBMap – Bushnell B. – sourceforge.net/projects/bbmap/). Raw reads were filtered for artifacts using BBDUK based on the following criteria: more than one N, quality scores with an average score less than 8 (before trimming), or reads shorter than 51 bp (after trimming). Reads were then mapped to masked versions of human, cat, and dog references and discarded if identity was greater than 95 % using BBMap. Reads were then masked using BBMask. Processed reads were assembled using SPAdes v3.6.2 [92] with the following parameters: –cov-cutoff auto –phred-offset 33 -t 8 - m 40 –careful -k 25,55,95 –12. Assembly contigs less than 1 kbp were discarded.

### Inoculating duckweed-bacteria samples

To study the bacterial colonization of duckweed, axenic duckweed was inoculated with the bacterial isolates described above. To inoculate duckweed with bacteria, a glycerol stock for the respective bacterium was used to inoculate a 5 mL liquid culture of LB or TSB and grown overnight at 28°C by shaking on a rotating platform at 220 rpm. A volume of 500 uL from the 5 mL liquid culture was then used to inoculate a 50 mL liquid culture of LB or TSB and grown overnight at 28°C at 220 rpm. The 50 mL bacterial culture was spun at 8000 rpm for 5 minutes at 4°C. The supernatant was then decanted, and the bacterial pellet was resuspended and washed with 0.5X SH. The sample was centrifuged as mentioned above. The resulting bacterial pellet was resuspended in 0.5X SH media and diluted to an OD600 of 0.2 in a final volume of 50 mL in a glass plant tissue culture vessel (Phytotechnology Laboratories, Catalog #C1770). Duckweed was then transferred to this 50 mL bacterial culture to cover the entire surface of the 50 mL bacterial culture. Inoculated duckweed was then incubated in a growth chamber under the same conditions used for duckweed propagation described above.

Wastewater samples were used to examine the colonization of duckweed by bacterial isolates in the presence of a microbial community. Wastewater samples were collected from the United Water Princeton Meadows wastewater treatment facility (Plainsboro, New Jersey, USA) after secondary clarification. For wastewater experiments, duckweed was inoculated as described above in 50 mL of non-sterile or filter-sterilized wastewater using 0.2 um polyethersulfone filters.

### Nucleic acid isolation from duckweed and bacteria

A bead-beating protocol was used to isolate nucleic acid from duckweed and bacteria. A combination of a 4 mm glass bead (OPS Diagnostics, Catalog #BAWG 4000-200-18), 0.5 grams of 1.7 mm zirconium beads (OPS Diagnostics, Catalog #BAWG 1700-300-22), and 0.5 grams of 100 um silica beads (OPS Diagnostics, Catalog #BAWG 100-200-10) was used for bead-beating to lyse samples. The lysis buffer consisted of 300 uL of high salt CTAB buffer (100 mM Tris-HCl pH 8, 2.0 M NaCl, 20 mM EDTA, 2 % CTAB) and 300 uL chloroform. All steps were carried out at room temperature. Duckweed, bacteria, or inoculated duckweed samples were transferred to bead-beating tubes with beads and lysis buffer then homogenized for 5 minutes (10 cycles of 30-second homogenization and 10-second pause) at 4000 rpm using an HT6 benchtop homogenizer from OPS Diagnostics (Lebanon, New Jersey, USA). Samples were then centrifuged at 16,000 X g for 5-10 minutes. Supernatants were transferred to new tubes and washed with 1X volume of 24:1 chloroform:isoamyl alcohol to remove protein precipitate and centrifuged at 16,000 X g for 5-10 minutes. This wash step was repeated. Supernatants were then transferred to new tubes and 0.5X volume of 7.5 M ammonium acetate and 2.5X volume of 95 % chilled ethanol were added [93]. Samples were centrifuged at 16,000 X g for 30 minutes to pellet the precipitated nucleic acid. The resulting sample pellets were washed with 70% chilled ethanol and centrifuged at 16,000 X g for 5-10 minutes. This step was repeated. Sample pellets were then air-dried and resuspended in 20 uL of sterile water or TE buffer. Nucleic acid concentration of samples were measured with a NanoDrop microvolume spectrophotometer (ThermoFisher Scientific).

### Detection of bacterial colonization by rDNA intergenic spacer analysis (RISA)

Primers for rRNA intergenic spacer analysis (RISA) were designed using previous reports [30,44,45,70] (File S2). The primers 16S-e1390f and 23S-e130r were selected to detect bacterial colonization of Lm5576 (File S2). RISA PCR reactions were prepared in a total volume of 25 uL consisting of: 0.5 mM MgCl2, 1X PCR buffer with Mg^2+^ (1.5 mM MgCl_2_, 10 mM KCl, 8 mM (NH_4_)_2_SO_4_, 10 mM Tris-HCl, pH 9.0, 0.05 % NP-40; Denville Scientific), 0.2 mM dNTPs, 0.8 uM forward primer, 0.8 uM reverse primer, and 2.5U of ChoiceTaq DNA polymerase (Denville Scientific, Catalog # CB4050-2). A volume of 2 uL of nucleic acids isolated from inoculated duckweed (∼100 ng/uL) or bacteria DNA (∼5 ng/uL) was added to RISA PCR reactions. RISA PCR reactions were executed using the following 3-stage thermocycler program: 1) denaturation stage of 95°C for 5 minutes, 2) cycling stage of 30 cycles consisting of 95°C for 15 seconds, 60°C for 30 seconds, 72°C for 1 minute 30 seconds, and 3) a final extension stage of 72°C for 5 minutes. RISA PCR products were visualized on a 1.0 % (w/v) agarose gel stained with ethidium bromide.

To verify sample quality and the relative amount of duckweed DNA in samples, primers were designed to the single copy, plant-specific *LEAFY* gene (File S2). *LEAFY* gene (*LFY*) primers were designed for dw9509 [47] and *L. minor* 5500 (Lm5500) [94]. Assembly and annotation files were retrieved from CoGe (https://genomevolution.org/coge/) for dw9509 (id 51364) and Lm5500 (id 27408). The LEAFY protein from Arabidopsis (NP_200993.1) was searched against the proteomes of dw9509 and Lm5500 using BLASTP v2.10.0+ [95]. Gene sequences were retrieved for top hits and a pairwise global alignment was generated using MUSCLE v3.8 [96]. The primers LmLFY-F and LmLFY-R were used to amplify the *LEAFY* gene from Lm5576 for endpoint PCR (*LmLFY* PCR)(File S2). The primers qLFY-F and qLFY-R were used to amplify the *LEAFY* gene from Lm5576 for real-time PCR (File S2). *LmLFY* PCR reactions were prepared in a total volume of 25 uL consisting of: 1X PCR buffer with Mg2+ (1.5 mM MgCl_2_, 10 mM KCl, 8 mM (NH_4_)_2_SO_4_, 10 mM Tris-HCl, pH 9.0, 0.05 % NP-40; Denville Scientific), 0.2 mM dNTPs, 0.4 uM forward primer, 0.4 uM reverse primer, 2U of ChoiceTaq DNA polymerase (Denville Scientific, Catalog # CB4050-2). A volume of 2 uL of nucleic acids isolated from inoculated duckweed (∼100 ng/uL) or duckweed DNA (∼5 ng/uL) was added to *LmLFY* PCR reactions. *LmLFY* PCR reactions were executed using the following 3-stage thermocycler program: 1) denaturation stage of 95°C for 5 minutes, 2) cycling stage of 28 cycles consisting of 95°C for 15 seconds, 60°C for 15 seconds, 72°C for 45 seconds, and 3) a final extension stage of 72°C for 5 minutes. *LmLFY* PCR products were visualized on a 1.0 % agarose gel stained with ethidium bromide.

### Confocal microscopy of Lm5576 colonized by bacteria

Lm5576 was inoculated with bacteria as described above. After seven days, inoculated Lm5576 tissue was harvested, washed with sterile H2O, and fixed in 1 mL of 4 % paraformaldehyde at RT in the dark overnight. The following day, the fixative solution was decanted and the fixed tissue was washed with 1 mL of 1X phosphate-buffered saline (PBS) twice. Fixed tissue was then stored at 4°C in 1 mL 1X PBS until further processing.

For confocal microscopy, paraformaldehyde-fixed Lm5576 plants were gently washed in 1X PBS and stained for DNA 16 hours at 4°C with SYBR Gold nucleic acid stain (ThermoFisher Scientific, Waltham, MA) diluted 1000X in 1X PBS. Samples were then washed with 1X PBS and stained with 0.5 mg/mL calcofluor white stain (Sigma-Aldrich, <u>St. Louis, MO</u>) for cellulose for 10 minutes at 22°C. Confocal images were acquired at 1 μm z-steps on a Zeiss LSM 710 (Carl Zeiss MicroImaging GmbH, Jena, Germany) scanning head confocal microscope with a Zeiss plan apo 40X/1.1 objective. Excitation lasers were 405 and 488 nm for the blue and green emission channels, respectively. The calcofluor white fluorescence was detected at 410-551 nm and the SYBR Gold fluorescence was detected at 533–572 nm. Laser dwell times were 2.55 μs for both channels. Image analysis (2D and 3D) was conducted using Zen (Zeiss) or Volocity (PerkinElmer, Waltham, MA).

### Strain-specific primer design and end-point PCR

Strain-specific primers were designed to detect the colonization of duckweed by specific bacterial strains (File S2). Two approaches were used to design primers, with both approaches requiring sequenced genomes of the respective bacterial strains. The first approach used Panseq v3.2.1 [97] to find unique sequences for primer design for endpoint PCR. The following configuration settings were used: minimumNovelRegionSize 500, novelRegionFinderMode unique, fragmentationSize 1000, percentIdentityCutoff 100, coreGenomeThreshold 2, runMode pan. The resulting unique sequences were then used for primer design. Primers were designed using the Primer3Plus web interface [98] with the following general settings: Primer Size Min 18, Primer Size Opt 20, Primer Size Max 25, Primer Tm Min 57, Primer Tm Opt 60, Primer Tm Max 63, Primer GC% Min 40, Primer GC% Opt 50, Primer GC% Max 60.

In the second approach, a custom computational pipeline composed of wrapper scripts (UniAmp) was implemented to find unique primers for each respective reference genome to use in real-time PCR. To accomplish this: 1) unique sequences to the reference genome were determined and 2) these unique sequences were used for primer design. To find unique reference genome sequences, first, query genomes were retrieved that were closely related to a reference genome. The Genome Taxonomy Database Toolkit (GTDB-tk) v1.7.0 was used to retrieve closely related query genomes from the Genome Taxonomy Database (GTDB) release 202) [99]. Additionally, the GenBank and RefSeq databases from the National Center for Biotechnology Information (NCBI) were remotely searched using the datasets v10.25.0 command line tool (https://github.com/ncbi/datasets). For this search, all genomes pertaining to the same genus as the reference genome were downloaded. RNAmmer v1.2 [100] was then used to extract the 16S rRNA gene sequences from these genomes. Only genomes whose 16S rRNA gene was > 97 % identical to the 16S rRNA gene from the reference genome were used as queries. Second, pairwise genome alignments were performed between each query genome and the reference genome using nucmer v3.1 [101]. Unique sequences in the reference genome, not found in any of the query genomes, were extracted. BedTools v2.25.0 [102] was used to find unique sequence intervals in the reference genome to build unique reference genome sequences. Only unique reference genome sequences that were 150-250 bp long and contained a GC content of 40-60 % were selected for further processing. As one last step to confirm sequences were unique to the reference genome, pairwise local alignments were performed between each unique sequence and query sequences from the same genus in the GenBank nucleotide database. Query sequences, from the same genus, were retrieved using the e-utilities from NCBI and compared using BLASTN v2.10.0+ [95]. Only the most unique reference sequence was used for primer design. To design primers, the unique reference sequence was used in a Primer-BLAST search using the specified parameters: PCR product size Min 100, PCR product size Max 200, # of primers to return 500, Database nr, Organism bacteria (taxid: 2), Primer must have at least 5 total mismatches to unintended targets, including at least 2 mismatches within the last 3 bps at the 3’ end, Primer Size Min 18, Primer Size Opt 22, Primer Size Max 26, Primer GC content (%) Min 40, Primer GC content (%) Max 60. Primer-BLAST results were saved as a HTML file and parsed using a custom Python script. In-silico PCR was then performed using USEARCH v11.0.667 [103] to determine the number of amplicons in the reference genome and in a selected set of query genomes. For each bacterial strain, primers with the fewest number of non-target amplicons found in the Primer-BLAST search, only 1 reference amplicon generated from in-silico PCR, and the lowest primer pair complementarity based on Primer-BLAST results were used for real-time PCR experiments. Primers were also subjected to PCR suitability tests using the PCR Primer Stats function of the online Sequence Manipulation Suite (https://www.bioinformatics.org/sms2/index.html) [104].

Strain-specific PCR reactions were prepared in a total volume of 25 uL consisting of 1X PCR buffer with Mg^2+^ (1.5 mM MgCl_2_, 10 mM KCl, 8 mM (NH_4_)_2_SO_4_, 10 mM Tris-HCl, pH 9.0, 0.05 % NP-40; Denville Scientific), 0.2 mM dNTPs, 0.4 uM forward primer, 0.4 uM reverse primer, and 2 Units of ChoiceTaq DNA polymerase (Denville Scientific, Catalog # CB4050-2). For Sp7 and DAB 1A specific PCR, 2% and 10 % DMSO were added respectively to end-point PCR reactions to avoid non-specific amplification. A volume of 2 uL from duckweed-bacteria nucleic acid samples (∼100 ng/uL) or bacteria DNA (5 ng/uL) was added to strain-specific PCR reactions. PCR reactions were executed using the following 3-stage thermocycler program: 1) a denaturation stage of 95°C for 5 minutes, 2) a cycling stage of 30 cycles consisting of 95°C for 15 seconds, 60°C for 15 seconds, 72°C for 30 seconds, and 3) a final extension stage of 72°C for 5 minutes. Strain-specific PCR products were visualized on a 1.0 % agarose gel stained with ethidium bromide.

### Quantification of bacterial colonization

Bacterial colonization of Lm5576 was quantified by real-time PCR (qPCR) using bacterial strain-specific and *qLFY* primers (File S2). For each sample, bacteria DNA and Lm5576 were quantified. Bacteria DNA was quantified using bacterial strain-specific primers designed by the custom UniAmp computational pipeline and Lm5576 DNA was quantified using *qLFY* primers complementary to the single-copy, plant-specific *LEAFY* gene. Bacteria DNA was divided by Lm5576 DNA for each sample to quantify bacterial colonization. For each qPCR reaction, a total volume of 20 uL was used and consisted of: 500 nM of forward primer, 500 nM of reverse primer, 1X Power SYBR Green PCR Master Mix (Thermo Fisher Scientific, Catalog # 4367659), and 5 uL of nucleic acid from inoculated duckweed or 5 uL of DNA from duckweed or bacteria alone. qPCR reactions were executed and analyzed using the StepOnePlus Real-Time PCR System from Applied Biosystems with StepOne software v2.2.2. The following settings were used: standard curve experiment, run method with a holding stage of 10 minutes at 95°C and cycling stage of 40 cycles consisting of 95°C for 15 seconds and 60°C for 1 minute. The following DNA standard ranges were used: 5, 0.5, 0.05, 0.005, 0.0005 ng/uL for bacteria DNA and 50, 5, 0.5, 0.05, 0.005 ng/uL for Lm5576 DNA. Standard curves generated for each primer set were successful if they met the following criteria: R2 > 0.97, efficiency between 80-110 %.

## Supplementary Materials

**Figure S1.**
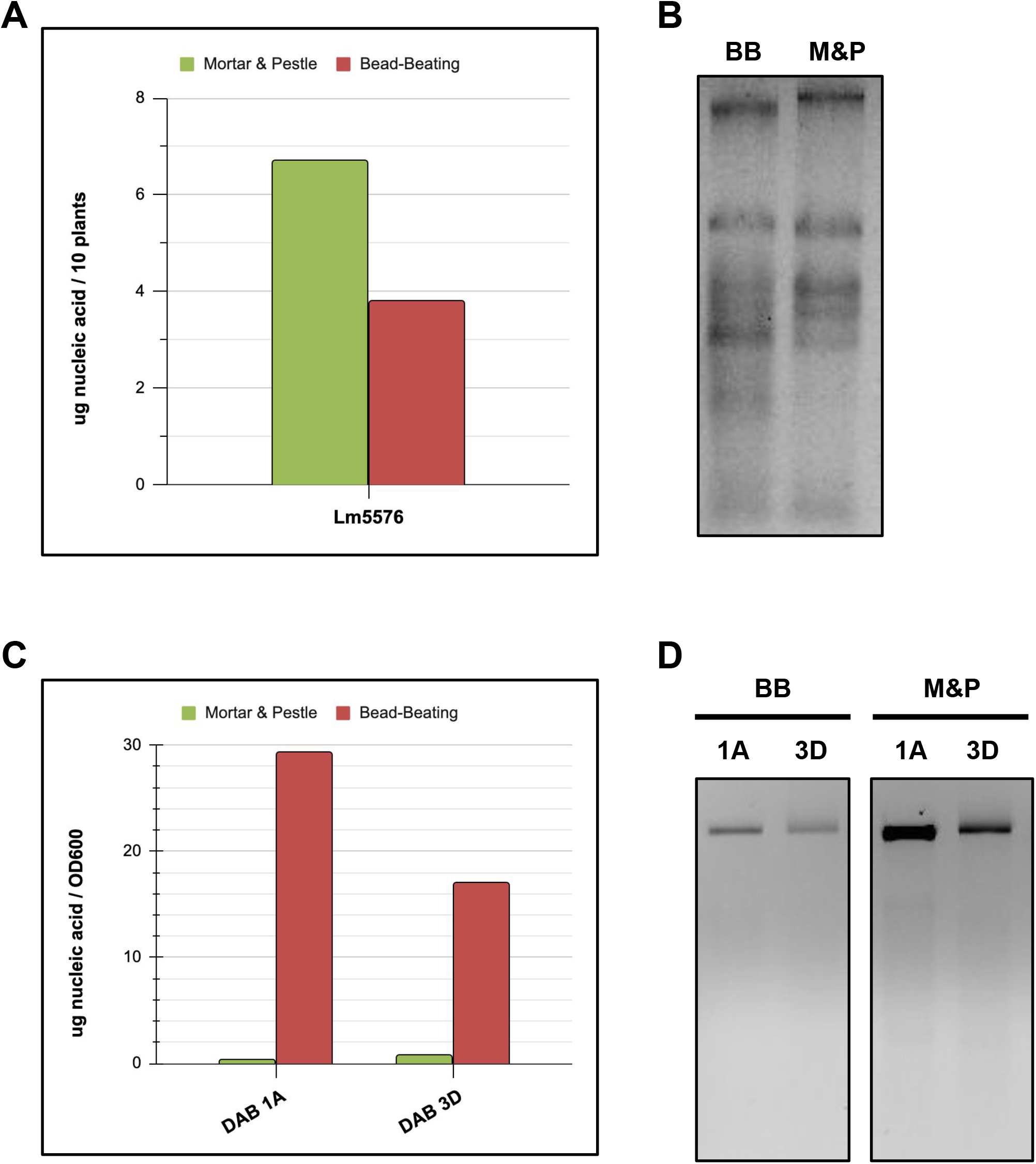
Nucleic acid isolation between mortar & pestle and bead-beating. Nucleic acid extraction was compared between mortar & pestle (M&P) and bead-beating (BB), using CTAB as the lysis buffer. **A)** Total micrograms (ug) of nucleic acids extracted per 10 plants of Lm5576 using bead-beating or mortar & pestle. To calculate the total ug of nucleic acid extracted, the nucleic acid concentration of the extract was multiplied by the total extract volume. **B)** Gel electrophoresis of approximately 500 nanograms of Lm5576 nucleic acids isolated with bead-beating or using mortar & pestle. **C)** Concentration of nucleic acids extracted from bacteria using bead-beating or mortar & pestle. The total micrograms (ug) of nucleic acid isolated was calculated by multiplying the nucleic acid concentration of extracts by the total extract volume. The total ug of nucleic acids isolated was then normalized to the optical density at 600 nm (OD600) of the liquid bacterial culture used for extraction. 1A = nucleic acids isolated from *Microbacterium sp*. RU370.1 (DAB 1A); 3D = nucleic acids isolated from *Bacillus sp*. RU9509.4 (DAB 3D). **D)** Gel electrophoresis of approximately 500 ng bacterial nucleic acids isolated with bead-beating or using mortar & pestle.

**Figure S2.**
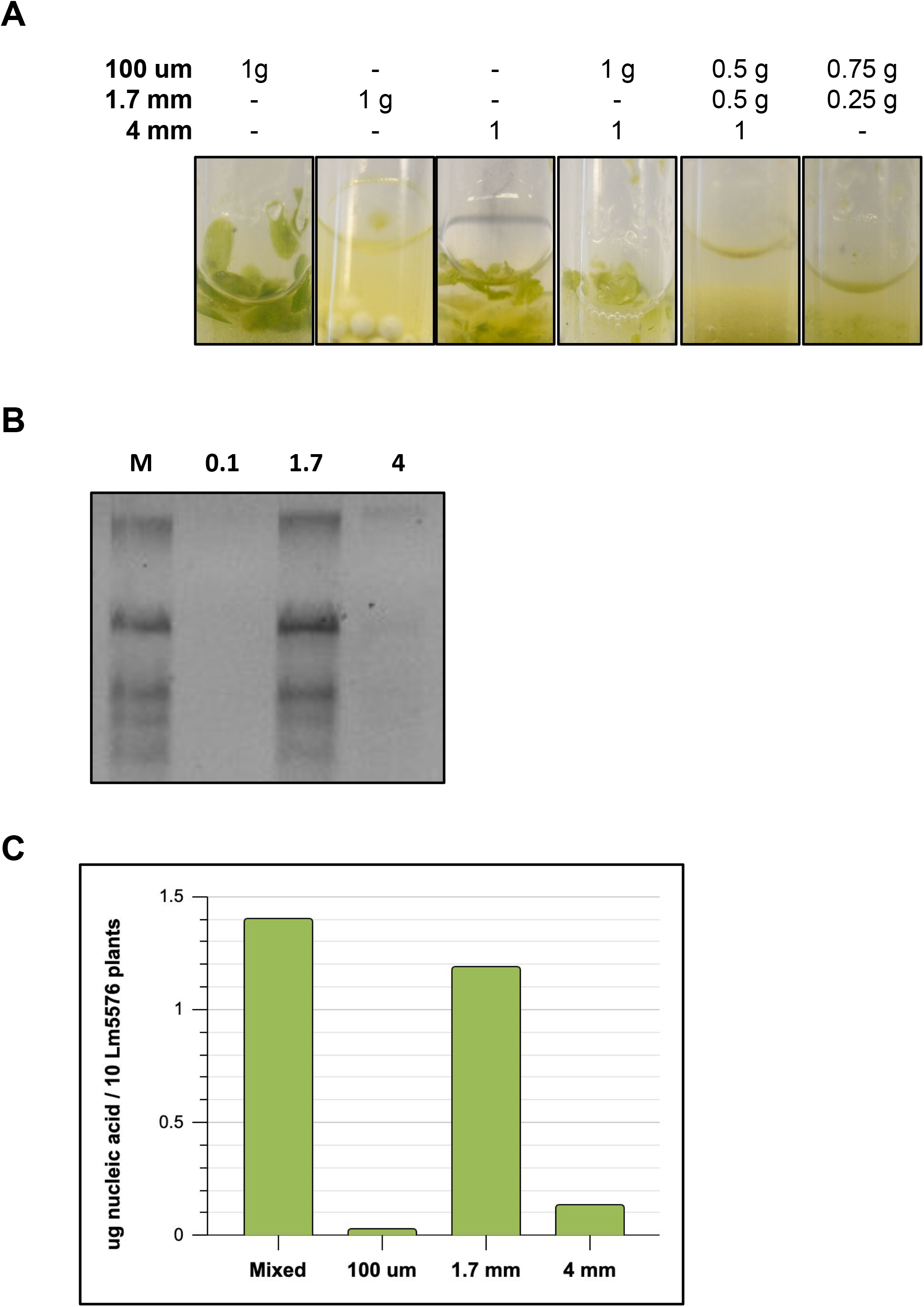
Nucleic acid isolation from Lm5576 with bead-beating. Different-sized beads were tested for their efficacy to extract nucleic acid from Lm5576. **A)** Homogenization of Lm5576 tissue by different bead sizes. **B)** Gel electrophoresis of approximately 500 ng nucleic acid isolated from Lm5576 using different bead sizes. M = Mixed; 0.1 = 100 um; 1.7 = 1.7 mm; 4 = 4 mm; Mixed = 0.5 g of 100 um beads, 0.5 g of 1.7 zirconium beads, and (1) 4 mm glass bead; Lm5576 = *Lemna minor* 5576. **C)** Total micrograms (ug) of nucleic acids extracted per 10 plants of Lm5576 using different bead sizes. To calculate the total ug of nucleic acid extracted, the nucleic acid concentration of the extract was multiplied by the total extract volume.

**Figure S3.**
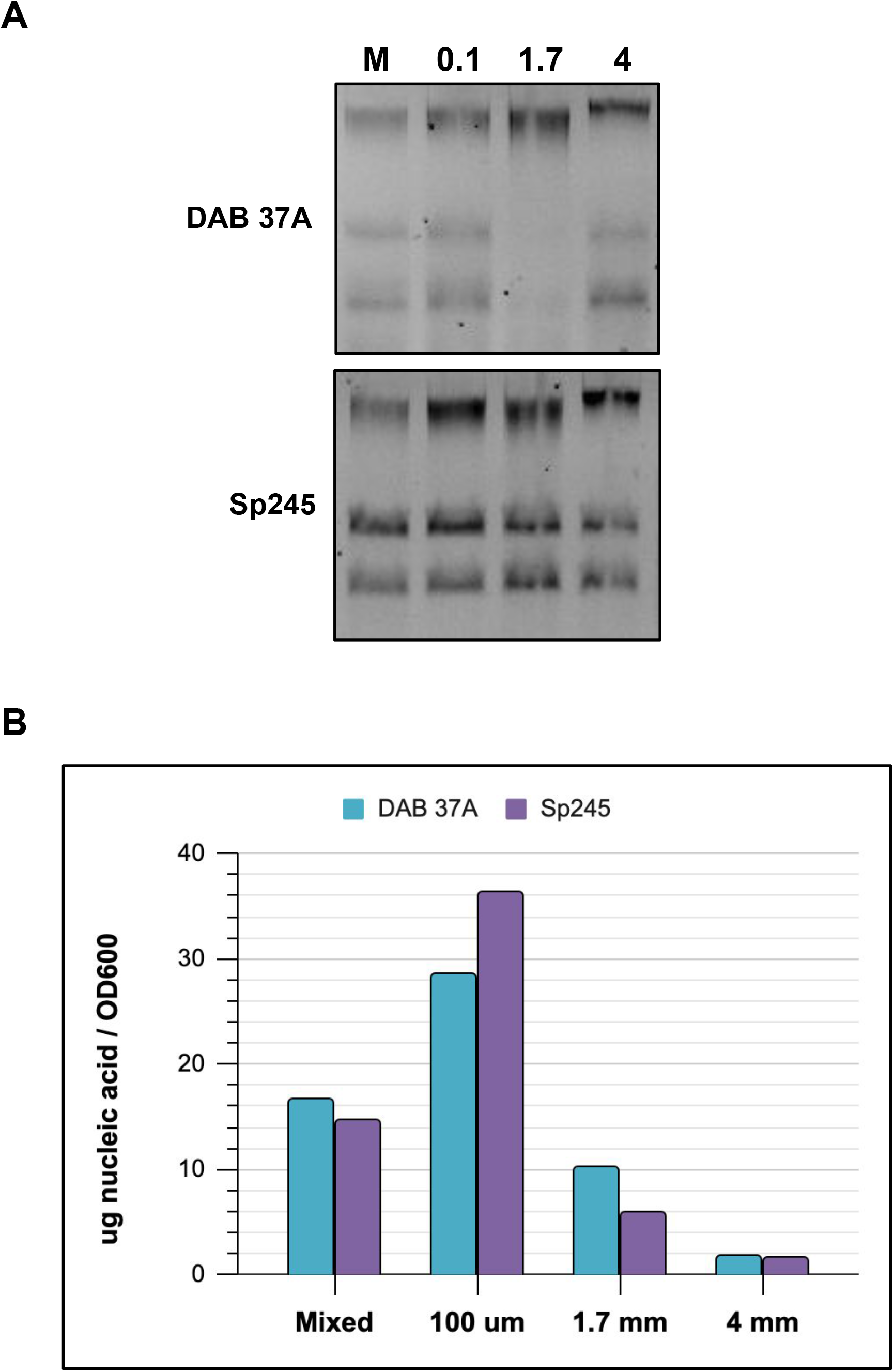
Nucleic acid isolation from bacteria with bead-beating. Different-sized beads were tested for extracting nucleic acid from bacteria. **A)** Gel electrophoresis of approximately 500 ng nucleic acid isolated from bacteria using different bead sizes. M = Mixed; 0.1 = 100 um; 1.7 = 1.7 mm; 4 = 4 mm; Mixed = 0.5 g of 100 um beads, 0.5 g of 1.7 zirconium beads, and (1) 4 mm glass bead; DAB 37A = nucleic acids isolated from DAB isolate 37A; Sp245 = nucleic acids isolated from *Azospirillum baldaniorum* Sp245. **B)** Concentration of nucleic acids extracted from bacteria using different bead sizes. The total micrograms (ug) of nucleic acid isolated was calculated by multiplying the nucleic acid concentration of extracts by the total extract volume. The total ug of nucleic acids isolated was then normalized to the optical density at 600 nm (OD600) of the liquid bacterial culture used for extraction.

**Figure S4.**
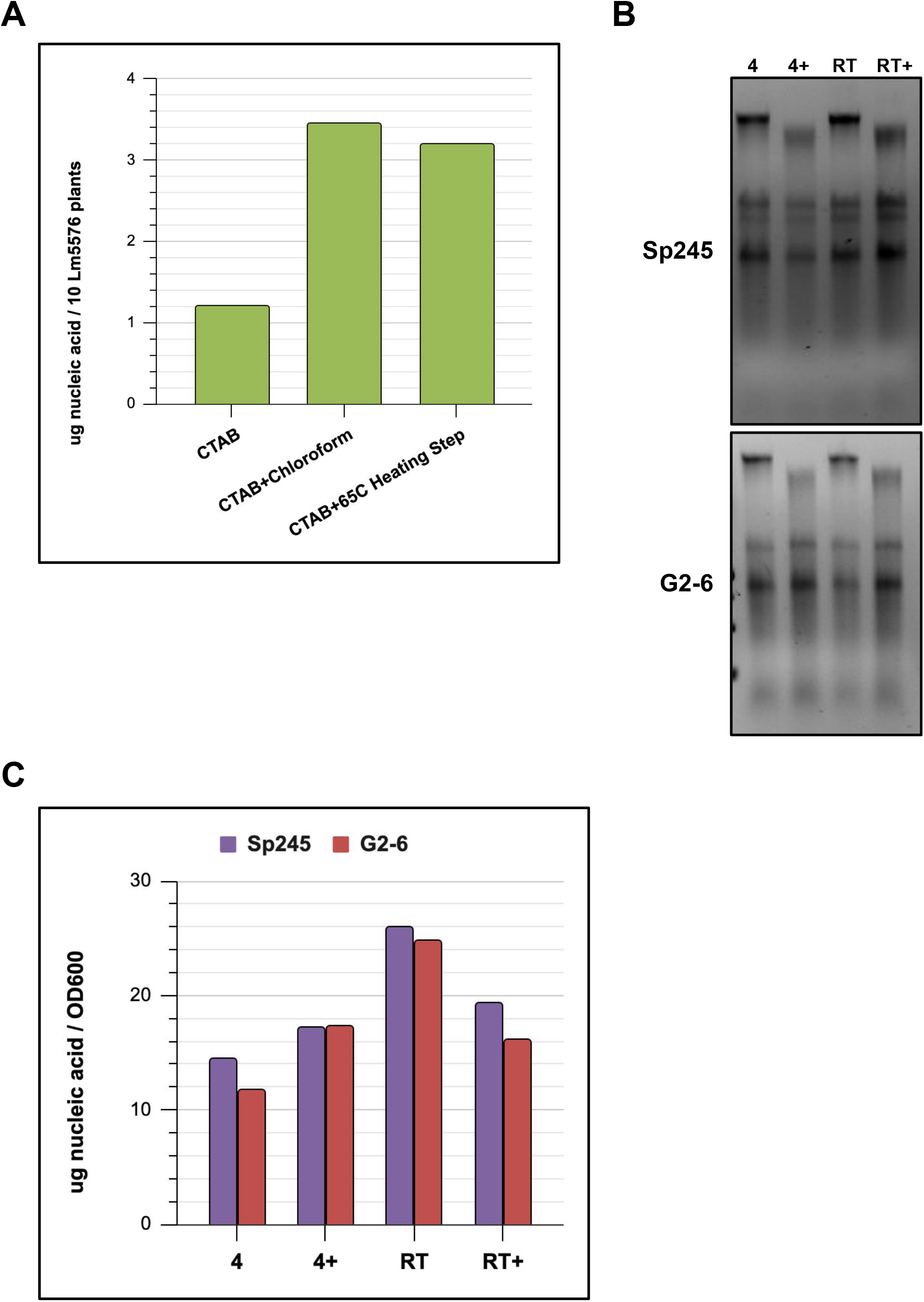
Optimization of nucleic acid extraction using a bead-beating protocol. Modifications to the lysis step of the bead-beating protocol were tested to improve nucleic acid extraction. **A)** Total micrograms (ug) of nucleic acids extracted per 10 plants of Lm5576 using different lysis modifications. To calculate the total ug of nucleic acid extracted, the nucleic acid concentration of the extract was multiplied by the total extract volume. CTAB = 600 uL CTAB lysis buffer; CTAB+Chloroform = 300 uL CTAB and 300 uL chloroform lysis buffer; CTAB+65°C Heating Step = 600 uL CTAB lysis buffer with 65°C heating step after lysis. **B)** Gel electrophoresis of approximately 500 ng nucleic acid isolated from bacteria using different lysis conditions. 4 = CTAB/chloroform lysis buffer and bead-beating at 4°C; 4+ = CTAB/chloroform lysis buffer plus 25 uL beta-mercaptoethanol and bead-beating at 4°C; RT = CTAB/chloroform lysis buffer and bead-beating at room temperature; RT+ = CTAB/chloroform lysis buffer plus 25 uL beta-mercaptoethanol and bead-beating at room temperature; Sp245 = nucleic acids isolated from *Azospirillum baldaniorum* Sp245; G2-6 = nucleic acids isolated from *Bacillus simplex* RUG2-6. **C)** Concentration of nucleic acids extracted from bacteria using different lysis conditions. The total micrograms (ug) of nucleic acid isolated was calculated by multiplying the nucleic acid concentration of extracts by the total extract volume. The total ug of nucleic acids isolated was then normalized to the optical density at 600 nm (OD600) of the liquid bacterial culture used for extraction.

**Figure S5.**
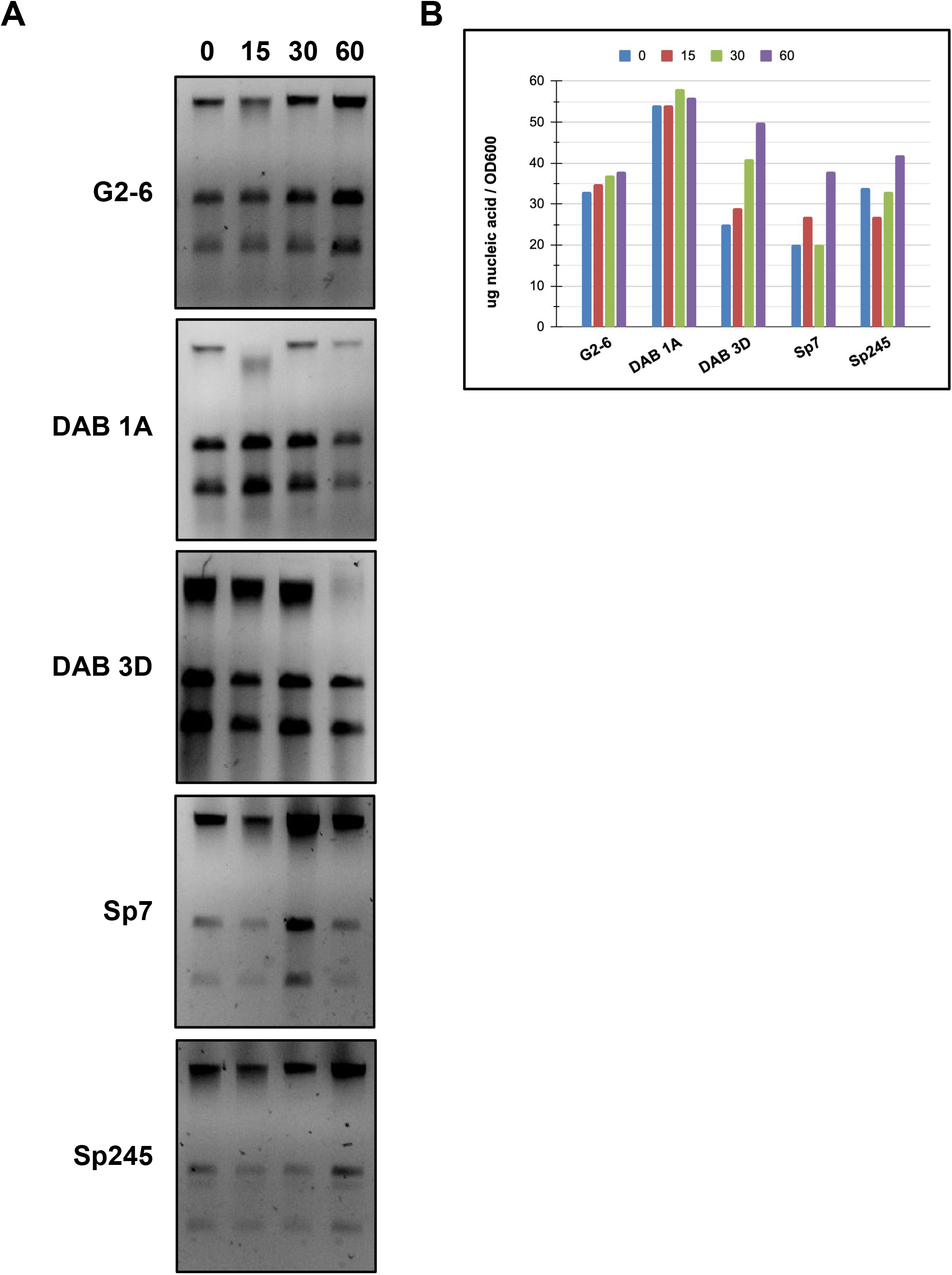
Nucleic acid isolation from different bacteria. Nucleic acids from different bacteria were extracted using the bead-beating protocol with different incubation times in the CTAB/chloroform lysis buffer. **A)** Gel electrophoresis of approximately 500 ng nucleic acids isolated from different bacteria using different incubation times in lysis buffer. 0 = no incubation in lysis buffer; 15 = 15-minute incubation in lysis buffer; 30 = 30-minute incubation in lysis buffer; 60 = 60-minute incubation in lysis buffer; G2-6 = nucleic acids isolated from *Bacillus simplex* RUG2-6; DAB 1A = nucleic acids isolated from *Microbacterium sp*. RU370.1; DAB 3D = nucleic acids isolated from *Bacillus sp*. RU9509.4; Sp7 = nucleic acids isolated from *Azospirillum brasilense* Sp7; Sp245 = nucleic acids isolated from *Azospirillum baldaniorum* Sp245. **B)** Concentration of nucleic acids extracted from bacteria using different incubation times in lysis buffer. The total micrograms (ug) of nucleic acid isolated was calculated by multiplying the nucleic acid concentration of extracts by the total extract volume. The total ug of nucleic acids isolated was then normalized to the optical density at 600 nm (OD600) of the liquid bacterial culture used for extraction.

**Figure S6.**
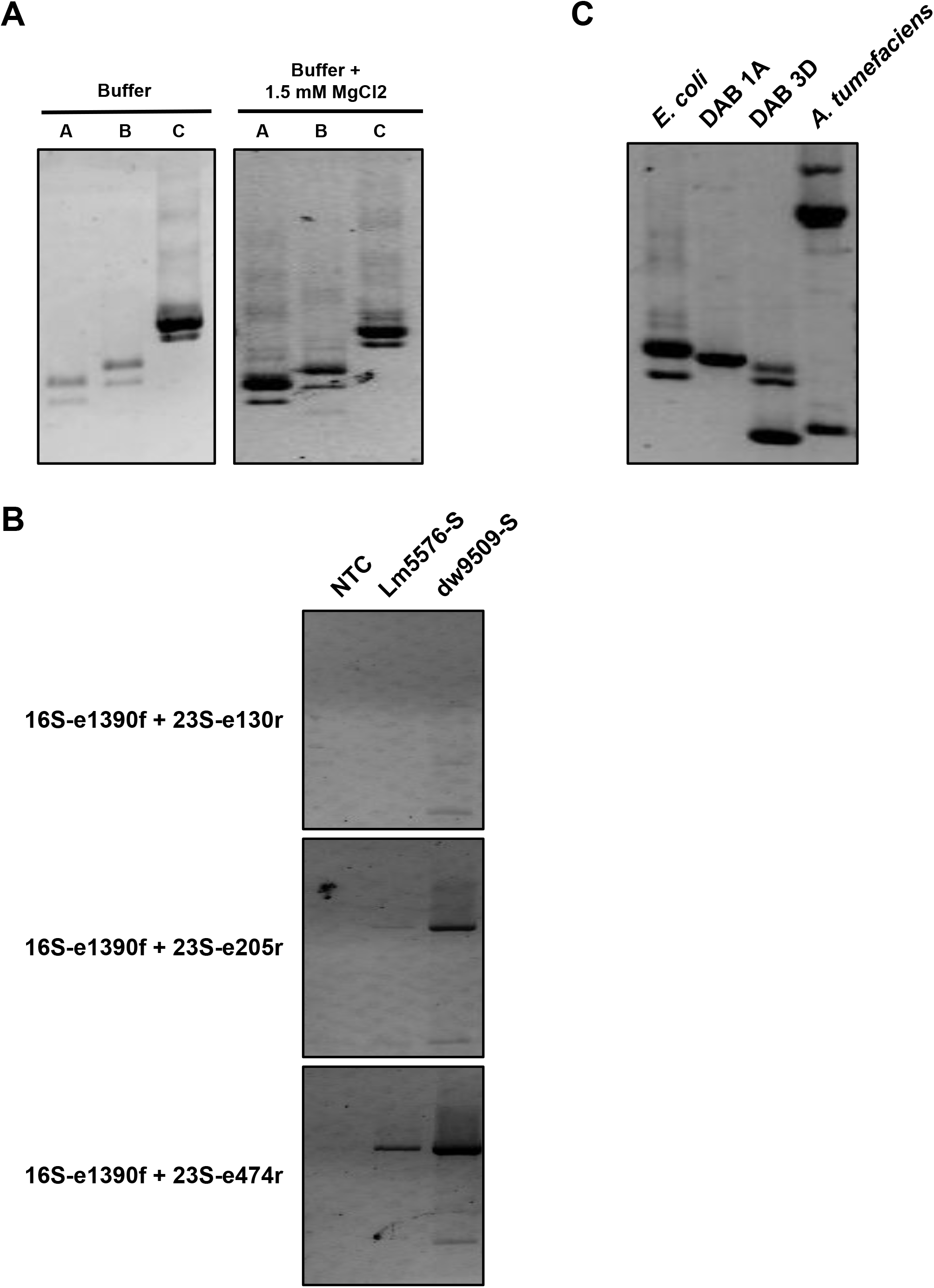
Amplification of bacteria DNA using RISA primers. **A)** Addition of magnesium chloride improves the amplification of bacteria DNA using RISA primers. Buffer = Choice Taq polymerase buffer (already contains 1.5 mM MgCl_2_); A = 16S-e1390f and 23S-e130r; B = 16S-e1390f and 23S-e205r; C = 16S-e1390f and 23S-e474r **B)** Different RISA primers were tested for their ability to amplify duckweed DNA. Lm5576-S = sterile *Lemna minor* 5576; dw9509-S = sterile *S. polyrhiza* 9509 **C)** RISA primers, 16S-e1390f and 23S-e130r, produce distinct fingerprints for different bacteria. NTC = no template control; *E. coli* = *Escherichia coli*; DAB 1A = *Microbacterium sp*. RU370.1; DAB 3D = *Bacillus sp*. RU9509.4; *A*.*tumefaciens* = *Agrobacterium tumefaciens*

**Figure S7.**
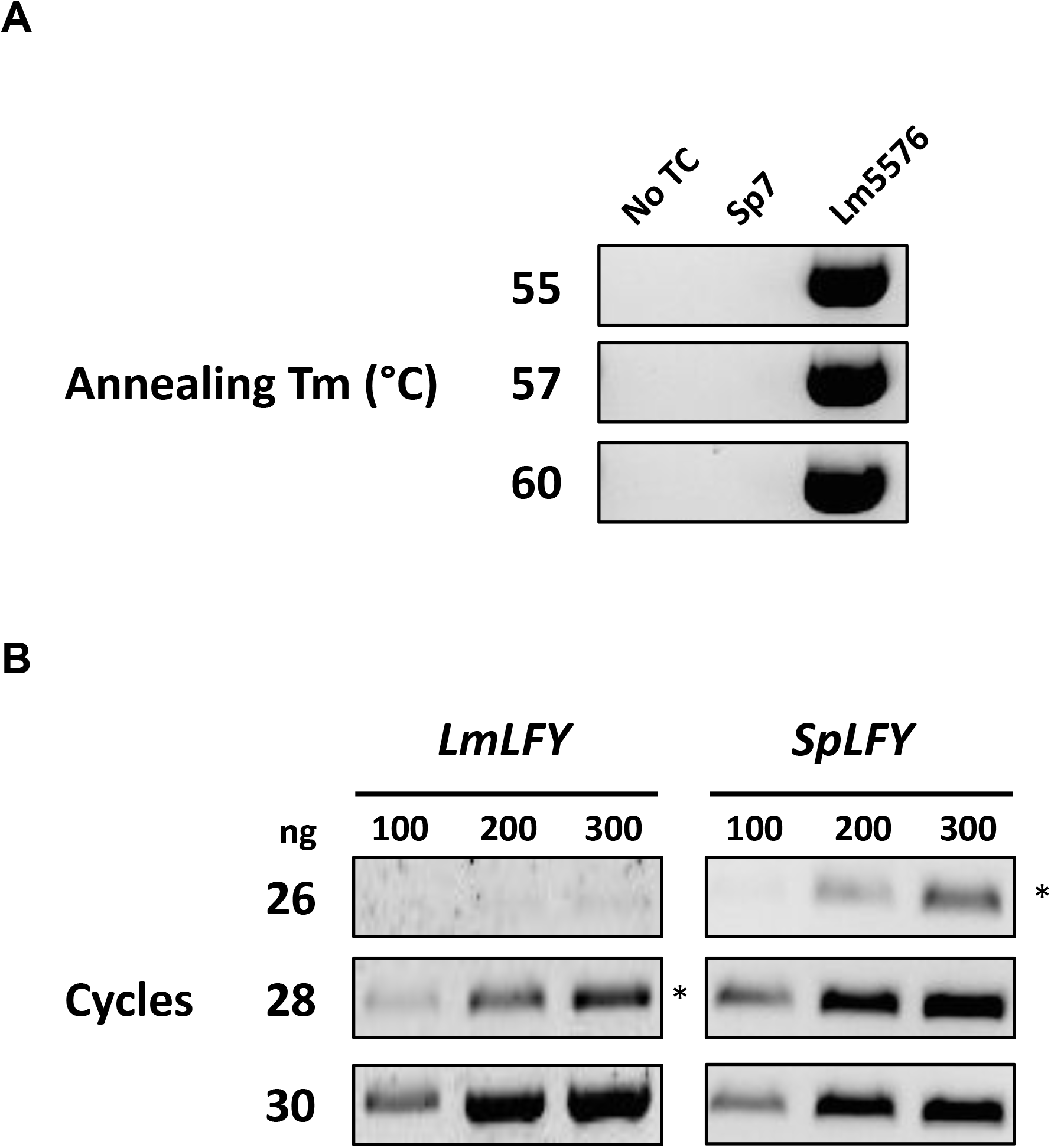
Optimization of *LEAFY* gene PCR. **A)** *LEAFY* gene PCR was performed on nucleic acids from bacteria and Lm5576 at different annealing temperatures. No TC = no template control; Sp7 = *Azospirillum brasilense* Sp7; Lm5576 = *Lemna minor* 5576 **B)** *LEAFY* gene PCR of Lm5576 and dw9509 nucleic acid at different concentrations using a different number of PCR cycles. * = the number of cycles selected for *LEAFY* gene PCR; *LmLFY* = PCR using LmLFY-F and LmLFY-R primers to amplify *LEAFY* gene from Lm5576; *SpLFY* = PCR using SpLFY-F and SpLFY-R primers to amplify *LEAFY* gene from dw5909

**Figure S8.**
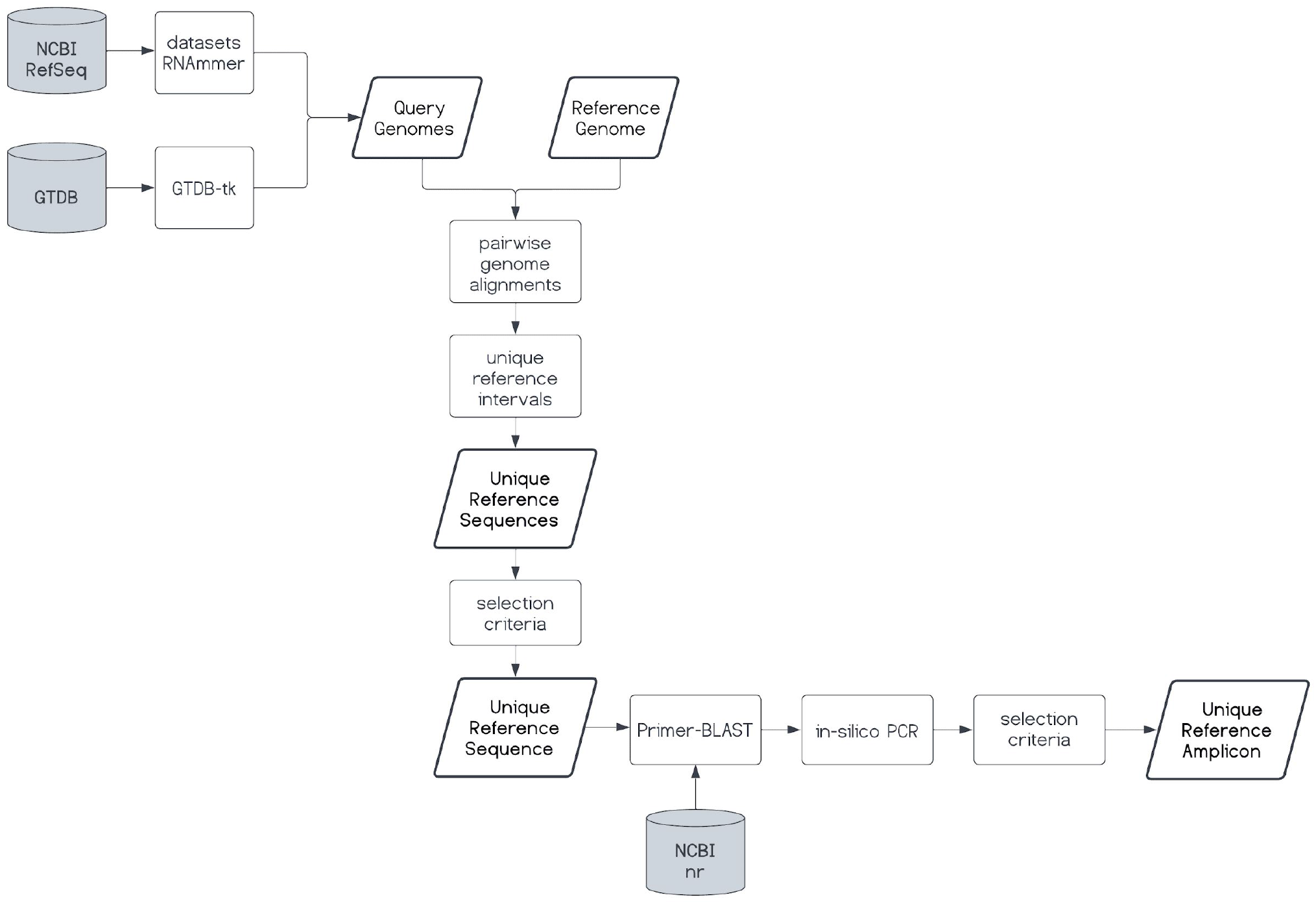
Overview of UniAmp computational pipeline to design strain-specific primers. The UniAmp pipeline can be conceptually split into 4 modules: 1) build a directory of query genomes, 2) retrieve unique sequences in a reference genome compared to query genomes, 3) select a unique reference sequence for primer design, and 4) design primers to the unique reference sequence.

**File S1. Metadata of bacterial isolates used in this study. A)** Isolation details, taxonomy, and colony morphology of bacterial isolates used in this study. Consensus 16S rRNA gene sequences were annotated with RDP classifier v.2.13 and 16S rRNA training set 18. **B)** Information on genomes generated in this study.

**File S2. Information on primers used in this study**.

**File S3. Design of duckweed *LEAFY* gene primers**. Pairwise alignment of *LEAFY* genes from *L. minor* 5500 and *S. polyrhiza* 9509. qLFY-F and qLFY-R = *LEAFY* gene primers used in real-time PCR to quantify *L. minor* and *S. polyrhiza* DNA; SpLFY-F and SpLFY-R = *LEAFY* gene primers specific to *S. polyrhiza* and used in end-point PCR; LmLFY-F and LmLFY-R = *LEAFY* gene primers specific to *L. minor* and used in end-point PCR

**File S4. Strain-specific primers generated using UniAmp computational pipeline**. Primer pairs highlighted in yellow were used in this study to detect the colonization of Lm5576 by G2-6, DAB 1A, Sp7, and Sp245 bacteria. Nontargets_organisms = number of non-targets amplified determined by Primer-BLAST, Organisms_amplified = number of organisms amplified determined by Primer-BLAST, For_pr_seq = forward primer sequence, Rev_pr_seq = reverse primer sequence, Self_complementarity = determined by Primer-BLAST, Self_3’_complementarity = determined by Primer-BLAST, Total_prpair_complementarity = sum of Self_complementarity and Self_3’_complementarity, Ref_amplicons = number of amplicons found in reference genome by UniAmp, Nonref_amplicons = number of amplicons found in selected query genomes by UniAmp, SMS_notes = manually curated notes from Sequence Manipulation Suite results

**File S5. 3D confocal microscopy of inoculated Lm5576 samples**. Calcofluor white was used to stain plant cellulose and visualized with the blue channel, SYBR Gold was used to stain DNA and visualized with the green channel, and chlorophyll autofluorescence was visualized with the red channel. Bacterial cells are stained green and are smaller in size compared to plant nuclei. White arrows point to cells of the respective bacterium in each sample.

**File S6. Attachment PCR results of confocal microscopy samples**.

## Data Availability Statement

Raw experimental data, bioinformatic analyses, and protocols used in this study can be found on figshare (https://figshare.com/account/home#/projects/155327). Protocols can be found on figshare ((https://figshare.com/account/home#/projects/155330) and protocols.io (https://protocols.io/workspaces/duckweed_microbiome). The UniAmp pipeline is available at https://github.com/kenscripts/UniAmp. Identifiers for 16S rRNA gene sequences and genomes generated in this study can be found in File S1. For the *Azospirillum brasilense* Sp7 genome, the JGI assembly with IMG genome id 2597490356 and GOLD analysis project ID Ga0060187 was used. For the *Azospirillum baldaniorum* Sp245 genome, the GenBank assembly GCA_000237365.1 was used.

## Author Contributions

Conceptualization: KA, EL, SL; Methodology: KA, EL, SS, WH, WC; Software: KA, TPM; Validation: KA, SS, WH, WC; Formal analysis: KA; Investigation: KA, SS, WC, WH, SG, TB; Resources: KA, EL, WC, SG, TB, SL; Data Curation: KA; Writing - Original: KA; Writing - Review & Editing: All authors; Visualization: KA, WC; Supervision: EL; Project administration: EL, SL; Funding acquisition: EL, SL

## Funding

Duckweed research at the Lam laboratory was supported in part by a grant from the Department of Energy (DE-SC0018244). The Lam lab was also supported by a Hatch project (#12116), and a Multi-State Capacity project (#NJ12710) from the New Jersey Agricultural Experiment Station at Rutgers University during this work. Contribution by the Facilities Integrating Collaborations for User Science (FICUS) initiative and under contract numbers DE732 AC02-05CH11231 (JGI) and DE-AC05-76RL01830 (EMSL) to the characterization of the duckweed microbiome is also gratefully acknowledged.

## Acknowledgments

A portion of this research was performed at the Environmental Molecular Sciences Laboratory, a national scientific user facility sponsored by the Department of Energy’s Office of Biological and Environmental Research and located at Pacific Northwest National Laboratory.

## Conflict of Interests

The authors declare no conflict of interest.

## Notes

### Competing Interest Statement

The authors have declared no competing interest.

## References

1. Acosta, K.; Appenroth, K.J.; Borisjuk, L.; Edelman, M.; Heinig, U.; Jansen, M.A.K.; Oyama, T.; Pasaribu, B.; Schubert, I.; Sorrels, S.; et al. Return of the Lemnaceae: Duckweed as a Model Plant System in the Genomics and Postgenomics Era. The Plant Cell 2021, 33, 3207–3234.

2. Fitzpatrick, C.R.; Salas-González, I.; Conway, J.M.; Finkel, O.M.; Gilbert, S.; Russ, D.; Teixeira, P.J.P.L.; Dangl, J.L. The Plant Microbiome: From Ecology to Reductionism and Beyond. Annu. Rev. Microbiol. 2020, 74, 81–100.

3. Acosta, K.; Xu, J.; Gilbert, S.; Denison, E.; Brinkman, T.; Lebeis, S.; Lam, E. Duckweed Hosts a Taxonomically Similar Bacterial Assemblage as the Terrestrial Leaf Microbiome. PLoS One 2020, 15, e0228560.

4. Xie, W.-Y.; Su, J.-Q.; Zhu, Y.-G. Phyllosphere Bacterial Community of Floating Macrophytes in Paddy Soil Environments as Revealed by Illumina High-Throughput Sequencing. Appl. Environ. Microbiol. 2015, 81, 522–532.

5. Inoue, D.; Hiroshima, N.; Ishizawa, H.; Ike, M. Whole Structures, Core Taxa, and Functional Properties of Duckweed Microbiomes. Bioresource Technology Reports 2022, 18, 101060.

6. Beilsmith, K.; Perisin, M.; Bergelson, J. Natural Bacterial Assemblages in Arabidopsis Thaliana Tissues Become More Distinguishable and Diverse during Host Development. MBio 2021, 12, doi:10.1128/mBio.02723-20.

7. Hacquard, S.; Garrido-Oter, R.; González, A.; Spaepen, S.; Ackermann, G.; Lebeis, S.; McHardy, A.C.; Dangl, J.L.; Knight, R.; Ley, R.; et al. Microbiota and Host Nutrition across Plant and Animal Kingdoms. Cell Host & Microbe 2015, 17, 603–616.

8. Wang, N.R.; Wiesmann, C.L.; Melnyk, R.A.; Hossain, S.S.; Chi, M.-H.; Martens, K.; Craven, K.; Haney, C.H. Commensal Pseudomonas Fluorescens Strains Protect Arabidopsis from Closely Related Pseudomonas Pathogens in a Colonization-Dependent Manner. mBio 2022, 13.

9. Innerebner, G.; Knief, C.; Vorholt, J.A. Protection of Arabidopsis Thaliana against Leaf-Pathogenic Pseudomonas Syringae by Sphingomonas Strains in a Controlled Model System. Appl. Environ. Microbiol. 2011, 77, 3202–3210.

10. Hassani, M.A.; Amine Hassani, M.; Durán, P.; Hacquard, S. Microbial Interactions within the Plant Holobiont. Microbiome 2018, 6.

11. Wippel, K.; Tao, K.; Niu, Y.; Zgadzaj, R.; Kiel, N.; Guan, R.; Dahms, E.; Zhang, P.; Jensen, D.B.; Logemann, E.; et al. Host Preference and Invasiveness of Commensal Bacteria in the Lotus and Arabidopsis Root Microbiota. Nat Microbiol 2021, 6, 1150–1162.

12. van Veen, J.A.; van Overbeek, L.S.; van Elsas, J.D. Fate and Activity of Microorganisms Introduced into Soil. Microbiol. Mol. Biol. Rev. 1997, 61, 121–135.

13. Gamalero, E.; Lingua, G.; Berta, G.; Lemanceau, P. Methods for Studying Root Colonization by Introduced Beneficial Bacteria. Sustainable Agriculture 2009, 601–615.

14. Katagiri, F.; Thilmony, R.; He, S.Y. The Arabidopsis Thaliana-Pseudomonas Syringae Interaction. arbo.j 2002, 2002, doi:10.1199/tab.0039.

15. Tornero, P.; Dangl, J.L. A High-Throughput Method for Quantifying Growth of Phytopathogenic Bacteria in Arabidopsis Thaliana. The Plant Journal 2002, 28, 475–481.

16. Haney, C.H.; Samuel, B.S.; Bush, J.; Ausubel, F.M. Associations with Rhizosphere Bacteria Can Confer an Adaptive Advantage to Plants. Nature Plants 2015, 1.

17. Zinniel, D.K.; Lambrecht, P.; Harris, N.B.; Feng, Z.; Kuczmarski, D.; Higley, P.; Ishimaru, C.A.; Arunakumari, A.; Barletta, R.G.; Vidaver, A.K. Isolation and Characterization of Endophytic Colonizing Bacteria from Agronomic Crops and Prairie Plants. Appl. Environ. Microbiol. 2002, 68, 2198–2208.

18. Niu, B.; Paulson, J.N.; Zheng, X.; Kolter, R. Simplified and Representative Bacterial Community of Maize Roots. Proc. Natl. Acad. Sci. U. S. A. 2017, 114, E2450–E2459.

19. O’Banion, B.S.; O’Neal, L.; Alexandre, G.; Lebeis, S.L. Bridging the Gap Between Single-Strain and Community-Level Plant-Microbe Chemical Interactions. Mol. Plant. Microbe. Interact. 2020, 33, 124–134.

20. Reinhold-Hurek, B.; Hurek, T. Life in Grasses: Diazotrophic Endophytes. Trends Microbiol. 1998, 6, 139–144.

21. James, E.K.; Gyaneshwar, P.; Mathan, N.; Barraquio, W.L.; Reddy, P.M.; Iannetta, P.P.M.; Olivares, F.L.; Ladha, J.K. Infection and Colonization of Rice Seedlings by the Plant Growth-Promoting Bacterium Herbaspirillum Seropedicae Z67. Mol. Plant. Microbe. Interact. 2002, 15, 894–906.

22. Schloter, M.; Hartmann, A. Endophytic and Surface Colonization of Wheat Roots (Triticum Aestivum) by Different Azospirillum Brasilense Strains Studied with Strain-Specific Monoclonal Antibodies. Symbiosis 1998, 25, 159–179.

23. Yamaga, F.; Washio, K.; Morikawa, M. Sustainable Biodegradation of Phenol by Acinetobacter Calcoaceticus P23 Isolated from the Rhizosphere of Duckweed Lemna Aoukikusa. Environ. Sci. Technol. 2010, 44, 6470–6474.

24. Assmus, B.; Hutzler, P.; Kirchhof, G.; Amann, R.; Lawrence, J.R.; Hartmann, A. In Situ Localization of Azospirillum Brasilense in the Rhizosphere of Wheat with Fluorescently Labeled, rRNA-Targeted Oligonucleotide Probes and Scanning Confocal Laser Microscopy. Appl. Environ. Microbiol. 1995, 61, 1013–1019.

25. Chelius, M.K.; Triplett, E.W. Immunolocalization of Dinitrogenase Reductase Produced by Klebsiella Pneumoniae in Association with Zea Mays L. Appl. Environ. Microbiol. 2000, 66, 783–787.

26. Hurek, T.; Reinhold-Hurek, B.; Van Montagu, M.; Kellenberger, E. Root Colonization and Systemic Spreading of Azoarcus Sp. Strain BH72 in Grasses. J. Bacteriol. 1994, 176, 1913–1923.

27. Fan, B.; Borriss, R.; Bleiss, W.; Wu, X. Gram-Positive Rhizobacterium Bacillus Amyloliquefaciens FZB42 Colonizes Three Types of Plants in Different Patterns. J. Microbiol. 2012, 50, 38–44.

28. Tringe, S.G.; Hugenholtz, P. A Renaissance for the Pioneering 16S rRNA Gene. Curr. Opin. Microbiol. 2008, 11, 442–446.

29. Thompson, L.R.; Sanders, J.G.; McDonald, D.; Amir, A.; Ladau, J.; Locey, K.J.; Prill, R.J.; Tripathi, A.; Gibbons, S.M.; Ackermann, G.; et al. A Communal Catalogue Reveals Earth’s Multiscale Microbial Diversity. Nature 2017, 551, 457–463.

30. Baker, G.C.; Smith, J.J.; Cowan, D.A. Review and Re-Analysis of Domain-Specific 16S Primers. J. Microbiol. Methods 2003, 55, 541–555.

31. Tkacz, A.; Hortala, M.; Poole, P.S. Absolute Quantitation of Microbiota Abundance in Environmental Samples. Microbiome 2018, 6.

32. Guo, X.; Zhang, X.; Qin, Y.; Liu, Y.-X.; Zhang, J.; Zhang, N.; Wu, K.; Qu, B.; He, Z.; Wang, X.; et al. Host-Associated Quantitative Abundance Profiling Reveals the Microbial Load Variation of Root Microbiome. Plant Commun 2020, 1, 100003.

33. Lundberg, D.S.; Pramoj Na Ayutthaya, P.; Strauß, A.; Shirsekar, G.; Lo, W.-S.; Lahaye, T.; Weigel, D. Host-Associated Microbe PCR (hamPCR) Enables Convenient Measurement of Both Microbial Load and Community Composition. Elife 2021, 10, doi:10.7554/eLife.66186.

34. Janda, J.M.; Abbott, S.L. 16S rRNA Gene Sequencing for Bacterial Identification in the Diagnostic Laboratory: Pluses, Perils, and Pitfalls. J. Clin. Microbiol. 2007, 45, 2761–2764.

35. Mignard, S.; Flandrois, J.P. 16S rRNA Sequencing in Routine Bacterial Identification: A 30-Month Experiment. J. Microbiol. Methods 2006, 67, 574–581.

36. Yang, B.; Wang, Y.; Qian, P.-Y. Sensitivity and Correlation of Hypervariable Regions in 16S rRNA Genes in Phylogenetic Analysis. BMC Bioinformatics 2016, 17, 135.

37. Pei, A.Y.; Oberdorf, W.E.; Nossa, C.W.; Agarwal, A.; Chokshi, P.; Gerz, E.A.; Jin, Z.; Lee, P.; Yang, L.; Poles, M.; et al. Diversity of 16S rRNA Genes within Individual Prokaryotic Genomes. Appl. Environ. Microbiol. 2010, 76, 3886–3897.

38. Maroniche, G.A.; García, J.E.; Salcedo, F.; Creus, C.M. Molecular Identification of Azospirillum Spp.: Limitations of 16S rRNA and Qualities of rpoD as Genetic Markers. Microbiol. Res. 2017, 195, 1–10.

39. Gilbert, S.; Xu, J.; Acosta, K.; Poulev, A.; Lebeis, S.; Lam, E. Bacterial Production of Indole Related Compounds Reveals Their Role in Association Between Duckweeds and Endophytes. Front Chem 2018, 6, 265.

40. Gilbert, S.; Poulev, A.; Chrisler, W.; Acosta, K.; Orr, G.; Lebeis, S.; Lam, E. Auxin-Producing Bacteria from Duckweeds Have Different Colonization Patterns and Effects on Plant Morphology. Plants 2022, 11, doi:10.3390/plants11060721.

41. Steenhoudt, O.; Vanderleyden, J. Azospirillum, a Free-Living Nitrogen-Fixing Bacterium Closely Associated with Grasses: Genetic, Biochemical and Ecological Aspects. FEMS Microbiol. Rev. 2000, 24, 487–506.

42. Ferreira, N. dos S.; dos Santos Ferreira, N.; Anna, F.H.S.; Reis, V.M.; Ambrosini, A.; Volpiano, C.G.; Rothballer, M.; Schwab, S.; Baura, V.A.; Balsanelli, E.; et al. Genome-Based Reclassification of Azospirillum Brasilense Sp245 as the Type Strain of Azospirillum Baldaniorum Sp. Nov. International Journal of Systematic and Evolutionary Microbiology 2020, 70, 6203–6212.

43. Borisjuk, N.; Chu, P.; Gutierrez, R.; Zhang, H.; Acosta, K.; Friesen, N.; Sree, K.S.; Garcia, C.; Appenroth, K.J.; Lam, E. Assessment, Validation and Deployment Strategy of a Two-Barcode Protocol for Facile Genotyping of Duckweed Species. Plant Biol. 2015, 17 Suppl 1, 42–49.

44. García-Martínez, J.; Acinas, S.G.; Antón, A.I.; Rodríguez-Valera, F. Use of the 16S--23S Ribosomal Genes Spacer Region in Studies of Prokaryotic Diversity. J. Microbiol. Methods 1999, 36, 55–64.

45. Gürtler, V.; Stanisich, V.A. New Approaches to Typing and Identification of Bacteria Using the 16S-23S rDNA Spacer Region. Microbiology 1996, 142 (Pt 1), 3–16.

46. Michael, T.P.; Bryant, D.; Gutierrez, R.; Borisjuk, N.; Chu, P.; Zhang, H.; Xia, J.; Zhou, J.; Peng, H.; El Baidouri, M.; et al. Comprehensive Definition of Genome Features in Spirodela Polyrhiza by High-Depth Physical Mapping and Short-Read DNA Sequencing Strategies. Plant J. 2017, 89, 617–635.

47. Hoang, P.N.T.; Michael, T.P.; Gilbert, S.; Chu, P.; Motley, S.T.; Appenroth, K.J.; Schubert, I.; Lam, E. Generating a High-Confidence Reference Genome Map of the Greater Duckweed by Integration of Cytogenomic, Optical Mapping, and Oxford Nanopore Technologies. Plant J. 2018, 96, 670–684.

48. Dennis, P.G.; Miller, A.J.; Hirsch, P.R. Are Root Exudates More Important than Other Sources of Rhizodeposits in Structuring Rhizosphere Bacterial Communities? FEMS Microbiol. Ecol. 2010, 72, 313–327.

49. Baker, J.H.; Orr, D.R. Distribution of Epiphytic Bacteria on Freshwater Plants. J. Ecol. 1986, 74, 155–165.

50. Borisjuk, N.; Peterson, A.A.; Lv, J.; Qu, G.; Luo, Q.; Shi, L.; Chen, G.; Kishchenko, O.; Zhou, Y.; Shi, J. Structural and Biochemical Properties of Duckweed Surface Cuticle. Front Chem 2018, 6, 317.

51. Ware, A.; Jones, D.H.; Flis, P.; Smith, K.; Kümpers, B.; Yant, L.; Atkinson, J.A.; Wells, D.M.; Bishopp, A. Duckweed Roots Are Dispensable and Are on a Trajectory toward Vestigiality. bioRxiv 2022, 2022.01.05.475062.

52. Chaintreuil, C.; Giraud, E.; Prin, Y.; Lorquin, J.; Bâ, A.; Gillis, M.; de Lajudie, P.; Dreyfus, B. Photosynthetic Bradyrhizobia Are Natural Endophytes of the African Wild Rice Oryza Breviligulata. Appl. Environ. Microbiol. 2000, 66, 5437–5447.

53. Gopalaswamy, G.; Kannaiyan, S.; O’Callaghan, K.J.; Davey, M.R.; Cocking, E.C. The Xylem of Rice (Oryza Sativa) Is Colonized byAzorhizobium Caulinodans. Proceedings of the Royal Society of London. Series B: Biological Sciences 2000, 267, 103–107.

54. Baldani, V.L.D.; de B. Alvarez, M.A.; Baldani, J.I.; Döbereiner, J. Establishment of Inoculated Azospirillum Spp. in the Rhizosphere and in Roots of Field Grown Wheat and Sorghum. Nitrogen Fixation with Non-Legumes 1986, 35–46.

55. Tarrand, J.J.; Krieg, N.R.; Döbereiner, J. A Taxonomic Study of the Spirillum Lipoferum Group, with Descriptions of a New Genus, Azospirillum Gen. Nov. and Two Species, Azospirillum Lipoferum (Beijerinck) Comb. Nov. and Azospirillum Brasilense Sp. Nov. Can. J. Microbiol. 1978, 24, 967–980.

56. Gafny, R.; Okon, Y.; Kapulnik, Y.; Fischer, M. Adsorption of Azospirillum Brasilense to Corn Roots. Soil Biology and Biochemistry 1986, 18, 69–75.

57. Mori, K.; Toyama, T.; Sei, K. Surfactants Degrading Activities in the Rhizosphere of Giant Duckweed (Spirodela Polyrhiza). Japanese Journal of Water Treatment Biology 2005, 41, 129–140.

58. Ishizawa, H.; Kuroda, M.; Inoue, K.; Inoue, D.; Morikawa, M.; Ike, M. Colonization and Competition Dynamics of Plant Growth-Promoting/Inhibiting Bacteria in the Phytosphere of the Duckweed Lemna Minor. Microbial Ecology 2019, 77, 440–450.

59. Haro, C.; Anguita-Maeso, M.; Metsis, M.; Navas-Cortés, J.A.; Landa, B.B. Evaluation of Established Methods for DNA Extraction and Primer Pairs Targeting 16S rRNA Gene for Bacterial Microbiota Profiling of Olive Xylem Sap. Front. Plant Sci. 2021, 12, 640829.

60. Henderson, G.; Cox, F.; Kittelmann, S.; Miri, V.H.; Zethof, M.; Noel, S.J.; Waghorn, G.C.; Janssen, P.H. Effect of DNA Extraction Methods and Sampling Techniques on the Apparent Structure of Cow and Sheep Rumen Microbial Communities. PLoS One 2013, 8, e74787.

61. Walker, A.W.; Martin, J.C.; Scott, P.; Parkhill, J.; Flint, H.J.; Scott, K.P. 16S rRNA Gene-Based Profiling of the Human Infant Gut Microbiota Is Strongly Influenced by Sample Processing and PCR Primer Choice. Microbiome 2015, 3, 26.

62. Maukonen, J.; Simões, C.; Saarela, M. The Currently Used Commercial DNA-Extraction Methods Give Different Results of Clostridial and Actinobacterial Populations Derived from Human Fecal Samples. FEMS Microbiol. Ecol. 2012, 79, 697–708.

63. Salonen, A.; Nikkilä, J.; Jalanka-Tuovinen, J.; Immonen, O.; Rajilić-Stojanović, M.; Kekkonen, R.A.; Palva, A.; de Vos, W.M. Comparative Analysis of Fecal DNA Extraction Methods with Phylogenetic Microarray: Effective Recovery of Bacterial and Archaeal DNA Using Mechanical Cell Lysis. J. Microbiol. Methods 2010, 81, 127–134.

64. Leite, G.M.; Magan, N.; Medina, Á. Comparison of Different Bead-Beating RNA Extraction Strategies: An Optimized Method for Filamentous Fungi. J. Microbiol. Methods 2012, 88, 413–418.

65. Miller, D.N.; Bryant, J.E.; Madsen, E.L.; Ghiorse, W.C. Evaluation and Optimization of DNA Extraction and Purification Procedures for Soil and Sediment Samples. Applied and Environmental Microbiology 1999, 65, 4715–4724.

66. Yuan, S.; Cohen, D.B.; Ravel, J.; Abdo, Z.; Forney, L.J. Evaluation of Methods for the Extraction and Purification of DNA from the Human Microbiome. PLoS ONE 2012, 7, e33865.

67. Pollock, J.; Glendinning, L.; Wisedchanwet, T.; Watson, M. The Madness of Microbiome: Attempting To Find Consensus “Best Practice” for 16S Microbiome Studies. Applied and Environmental Microbiology 2018, 84.

68. Fouhy, F.; Clooney, A.G.; Stanton, C.; Claesson, M.J.; Cotter, P.D. 16S rRNA Gene Sequencing of Mock Microbial Populations-Impact of DNA Extraction Method, Primer Choice and Sequencing Platform. BMC Microbiol. 2016, 16, 123.

69. Brooks, J.P.; Edwards, D.J.; Harwich, M.D., Jr; Rivera, M.C.; Fettweis, J.M.; Serrano, M.G.; Reris, R.A.; Sheth, N.U.; Huang, B.; Girerd, P.; et al. The Truth about Metagenomics: Quantifying and Counteracting Bias in 16S rRNA Studies. BMC Microbiol. 2015, 15, 66.

70. Fisher, M.M.; Triplett, E.W. Automated Approach for Ribosomal Intergenic Spacer Analysis of Microbial Diversity and Its Application to Freshwater Bacterial Communities. Applied and Environmental Microbiology 1999, 65, 4630–4636.

71. Bodenhausen, N.; Bortfeld-Miller, M.; Ackermann, M.; Vorholt, J.A. A Synthetic Community Approach Reveals Plant Genotypes Affecting the Phyllosphere Microbiota. PLoS Genet. 2014, 10, e1004283.

72. Bowker, D.W.; Duffield, A.N.; Denny, P. Methods for the Isolation, Sterilization and Cultivation of Lemnaceae. Freshwater Biology 1980, 10, 385–388.

73. Huang, W.; Gilbert, S.; Poulev, A.; Acosta, K.; Lebeis, S.; Long, C.; Lam, E. Host-Specific and Tissue-Dependent Orchestration of Microbiome Community Structure in Traditional Rice Paddy Ecosystems. Plant and Soil 2020, 452, 379–395.

74. Bodenhausen, N.; Deslandes-Hérold, G.; Waelchli, J.; Held, A.; van der Heijden, M.G.A.; Schlaeppi, K. Relative qPCR to Quantify Colonization of Plant Roots by Arbuscular Mycorrhizal Fungi. Mycorrhiza 2021, 31, 137–148.

75. Garrido-Oter, R.; Nakano, R.T.; Dombrowski, N.; Ma, K.-W.; AgBiome Team McHardy, A.C.; Schulze-Lefert, P. Modular Traits of the Rhizobiales Root Microbiota and Their Evolutionary Relationship with Symbiotic Rhizobia. Cell Host Microbe 2018, 24, 155–167.e5.

76. Müller, D.B.; Vogel, C.; Bai, Y.; Vorholt, J.A. The Plant Microbiota: Systems-Level Insights and Perspectives. Annual Review of Genetics 2016, 50, 211–234.

77. Trivedi, P.; Leach, J.E.; Tringe, S.G.; Sa, T.; Singh, B.K. Plant–microbiome Interactions: From Community Assembly to Plant Health. Nature Reviews Microbiology 2020, 18, 607– 621.

78. Levy, A.; Salas Gonzalez, I.; Mittelviefhaus, M.; Clingenpeel, S.; Herrera Paredes, S.; Miao, J.; Wang, K.; Devescovi, G.; Stillman, K.; Monteiro, F.; et al. Genomic Features of Bacterial Adaptation to Plants. Nat. Genet. 2017, 50, 138–150.

79. Hardoim, P.R.; van Overbeek, L.S.; van Elsas, J.D. Properties of Bacterial Endophytes and Their Proposed Role in Plant Growth. Trends Microbiol. 2008, 16, 463–471.

80. Cole, B.J.; Feltcher, M.E.; Waters, R.J.; Wetmore, K.M.; Mucyn, T.S.; Ryan, E.M.; Wang, G.; Ul-Hasan, S.; McDonald, M.; Yoshikuni, Y.; et al. Genome-Wide Identification of Bacterial Plant Colonization Genes. PLoS Biol. 2017, 15, e2002860.

81. Ishizawa, H.; Kuroda, M.; Inoue, D.; Ike, M. Genome-Wide Identification of Bacterial Colonization and Fitness Determinants on the Floating Macrophyte, Duckweed. Commun Biol 2022, 5, 68.

82. Duca, D.; Lorv, J.; Patten, C.L.; Rose, D.; Glick, B.R. Indole-3-Acetic Acid in Plant–microbe Interactions. Antonie van Leeuwenhoek 2014, 106, 85–125.

83. Tzipilevich, E.; Russ, D.; Dangl, J.L.; Benfey, P.N. Plant Immune System Activation Is Necessary for Efficient Root Colonization by Auxin-Secreting Beneficial Bacteria. Cell Host Microbe 2021, 29, 1507–1520.e4..

84. Ishizawa, H.; Kuroda, M.; Morikawa, M.; Ike, M. Evaluation of Environmental Bacterial Communities as a Factor Affecting the Growth of Duckweed Lemna Minor. Biotechnology for Biofuels 2017, 10.

85. Ishizawa, H.; Kuroda, M.; Inoue, D.; Morikawa, M.; Ike, M. Community Dynamics of Duckweed-Associated Bacteria upon Inoculation of Plant Growth-Promoting Bacteria. FEMS Microbiol. Ecol. 2020, 96, doi:10.1093/femsec/fiaa101.

86. Vorholt, J.A.; Vogel, C.; Carlström, C.I.; Müller, D.B. Establishing Causality: Opportunities of Synthetic Communities for Plant Microbiome Research. Cell Host Microbe 2017, 22, 142–155.

87. Carper, D.L.; Lawrence, T.J.; Carrell, A.A.; Pelletier, D.A.; Weston, D.J. DISCo-Microbe: Design of an Identifiable Synthetic Community of Microbes. PeerJ 2020, 8, e8534.

88. Miché, L.; Balandreau, J. Effects of Rice Seed Surface Sterilization with Hypochlorite on Inoculated Burkholderia Vietnamiensis. Appl. Environ. Microbiol. 2001, 67, 3046–3052.

89. Wang, Q.; Garrity, G.M.; Tiedje, J.M.; Cole, J.R. Naive Bayesian Classifier for Rapid Assignment of rRNA Sequences into the New Bacterial Taxonomy. Appl. Environ. Microbiol. 2007, 73, 5261–5267.

90. Zerbino, D.R.; Birney, E. Velvet: Algorithms for de Novo Short Read Assembly Using de Bruijn Graphs. Genome Research 2008, 18, 821–829.

91. Butler, J.; MacCallum, I.; Kleber, M.; Shlyakhter, I.A.; Belmonte, M.K.; Lander, E.S.; Nusbaum, C.; Jaffe, D.B. ALLPATHS: De Novo Assembly of Whole-Genome Shotgun Microreads. Genome Res. 2008, 18, 810–820.

92. Bankevich, A.; Nurk, S.; Antipov, D.; Gurevich, A.A.; Dvorkin, M.; Kulikov, A.S.; Lesin, V.M.; Nikolenko, S.I.; Pham, S.; Prjibelski, A.D.; et al. SPAdes: A New Genome Assembly Algorithm and Its Applications to Single-Cell Sequencing. J. Comput. Biol. 2012, 19, 455– 477.

93. Crouse, J.; Amorese, D. Ethanol Precipitation: Ammonium Acetate as an Alternative to Sodium Acetate. Focus 1996, 19, 17–20.

94. Van Hoeck, A.; Horemans, N.; Monsieurs, P.; Cao, H.X.; Vandenhove, H.; Blust, R. The First Draft Genome of the Aquatic Model Plant Lemna Minor Opens the Route for Future Stress Physiology Research and Biotechnological Applications. Biotechnol. Biofuels 2015, 8, 188.

95. Camacho, C.; Coulouris, G.; Avagyan, V.; Ma, N.; Papadopoulos, J.; Bealer, K.; Madden, T.L. BLAST : Architecture and Applications. BMC Bioinformatics 2009, 10.

96. Edgar, R.C. MUSCLE: Multiple Sequence Alignment with High Accuracy and High Throughput. Nucleic Acids Res. 2004, 32, 1792–1797.

97. Laing, C.; Buchanan, C.; Taboada, E.N.; Zhang, Y.; Kropinski, A.; Villegas, A.; Thomas, J.E.; Gannon, V.P.J. Pan-Genome Sequence Analysis Using Panseq: An Online Tool for the Rapid Analysis of Core and Accessory Genomic Regions. BMC Bioinformatics 2010, 11, 461.

98. Untergasser, A.; Nijveen, H.; Rao, X.; Bisseling, T.; Geurts, R.; Leunissen, J.A.M. Primer3Plus, an Enhanced Web Interface to Primer3. Nucleic Acids Res. 2007, 35, W71– W74.

99. Chaumeil, P.-A.; Mussig, A.J.; Hugenholtz, P.; Parks, D.H. GTDB-Tk: A Toolkit to Classify Genomes with the Genome Taxonomy Database. Bioinformatics 2019, 36, 1925–1927.

100. Lagesen, K.; Hallin, P.; Rødland, E.A.; Staerfeldt, H.-H.; Rognes, T.; Ussery, D.W. RNAmmer: Consistent and Rapid Annotation of Ribosomal RNA Genes. Nucleic Acids Res. 2007, 35, 3100–3108.

101. Kurtz, S.; Phillippy, A.; Delcher, A.L.; Smoot, M.; Shumway, M.; Antonescu, C.; Salzberg, S.L. Versatile and Open Software for Comparing Large Genomes. Genome Biol. 2004, 5, R12.

102. Quinlan, A.R.; Hall, I.M. BEDTools: A Flexible Suite of Utilities for Comparing Genomic Features. Bioinformatics 2010, 26, 841–842.

103. Edgar, R.C. Search and Clustering Orders of Magnitude Faster than BLAST. Bioinformatics 2010, 26, 2460–2461.

104. Stothard, P. The Sequence Manipulation Suite: JavaScript Programs for Analyzing and Formatting Protein and DNA Sequences. Biotechniques 2000, 28, 1102, 1104.

